# DeepCryoRNA: deep learning-based RNA structure reconstruction from cryo-EM maps

**DOI:** 10.1101/2025.04.05.647396

**Authors:** Jun Li, Shi-Jie Chen

## Abstract

RNA molecules play a crucial role in various cellular functions, making understanding their three-dimensional (3D) structures vitally important. Cryogenic electron microscopy (cryo-EM) has significantly advanced the study of RNA structures. However, most existing modeling tools are primarily developed for proteins, making them less effective for RNAs due to the unique structural variability and flexibility of an RNA. Addressing the limitations of current cryo-EM based RNA structure determination, we develop DeepCryoRNA, a novel deep learning approach tailored for automated reconstruction of RNA 3D structures from cryo-EM maps at high to medium resolutions. The approach was rigorously evaluated using a benchmark set of 51 RNA structures, resolved by cryo-EM at resolutions finer than 6.0 Å. The results demonstrate DeepCryoRNA’s marked accuracy over existing methods. The methodology of DeepCryoRNA stands out for its accurate atomistic predictions and advanced global sequence alignment capabilities, enabling more accurate sequence assignments. These features collectively establish DeepCryoRNA as a highly efficient and effective tool for RNA 3D structure reconstruction from cryo-EM maps, advancing the field of RNA structural biology.

## 1 Introduction

RNA is increasingly recognized as a crucial regulator in various cellular processes beyond its conventional role in protein synthesis [1, 2]. In recent years, structure-based understanding of multifaceted functions of RNA has paved the way for significant advancements in RNA-based therapies [3, 4]. A groundbreaking example of the progress is the development of mRNA vaccines [5], a revolutionary RNA-based therapeutic strategy that has helped mitigate the COVID-19 pandemic.

The therapeutic potential of RNA underscores the critical importance of further understanding RNA 3D structures. To accurately determine these structures, a range of experimental techniques, including conventional X-ray crystallography, nuclear magnetic resonance (NMR) spectroscopy, and the novel cryo-EM, have led to significant success. Cryo-EM, in particular, has emerged as a revolutionary tool [6] for its power to capture dynamic structures and resolve large structures without crystallization. The effectiveness of cryo-EM, however, relies on specialized software capable of modeling structures from cryo-EM maps. The majority of the available advanced modeling tools are primarily tailored for protein structures [7–20], and their applicability to RNA structure modeling is quite limited. Moreover, RNA structures exhibit a higher degree of flexibility and diversity than their protein counterparts, making the construction of 3D conformations from cryo-EM particularly challenging [21, 22].

Currently, only a few approaches are available for automated modeling of RNA structures from cryo-EM maps. These methods include auto-DRRAFTER [23], and the recent deep learning-based approaches, DeepTracer-2.0 [24] (referred to as “Deep-Tracer” in the current study) and CryoREAD [25]. In addition to cryo-EM maps, auto-DRRAFTER [23] depends also on accurate RNA secondary structures and entails both CPU- and time-consuming conformational sampling to iteratively identify the converged structures. DeepTracer [24] exhibits the capability to concurrently or separately reconstruct 3D structures of proteins and nucleic acids including DNA and RNA. When focusing on nucleic acid structure prediction, it leverages a trained nucleotide U-Net neural network [26, 27] to predict the positions of three critical atom types, namely P, C1’, and C4’. Complementing this, a dedicated backbone U-Net neural network is deployed to predict the sugar-phosphate backbones and the nitrogenous bases. Subsequently, a nucleotide post-processing phase is implemented to ensure the reconstruction of all-atom structures. CryoREAD [25] specializes in the reconstruction of 3D structures for nucleic acids (DNA and RNA) from cryo-EM maps, regardless of the presence of proteins. It uses two stages of U-Net neural networks [26, 27] to predict the positions of sugar, phosphate, and four types of bases. Subsequently, it proceeds to trace the backbone structure and assigns the RNA sequence via fragment sequence alignment. Ultimately, CryoREAD finalizes the reconstruction process by producing detailed all-atom structures. Unlike auto-DRRAFTER [23], the two deep learning-based approaches, DeepTracer [24] and CryoREAD [25], stand out for their full automation and efficiency in modeling RNA structures from cryo-EM maps.

In order to assess the practical performance of the two deep learning-based methods, DeepTracer [24] and CryoREAD [25], in reconstructing RNA structures from RNA cryo-EM maps, we conducted a comprehensive benchmark test. This evaluation covered all 51 available pure RNA structures solved via cryo-EM, each with a cryo-EM map resolution better than 6.0 Å. The raw cryo-EM maps downloaded from the Electron Microscopy Data Bank (EMDB) [28] and the corresponding RNA sequences were submitted to the DeepTracer and CryoREAD servers. The native RNA structures were downloaded from the RCSB Protein Data Bank (PDB) [29] for comparison. However, the predicted 3D structures, which showed large root mean square deviations (RMSDs) compared to the native RNA structures, are fall short of satisfaction. Specifically, in the majority of the test cases, DeepTracer [24] typically predicted only a few nucleotides or phosphorus atoms. CryoREAD [25] effectively generated the expected number of nucleotides from the cryo-EM maps. However, at the chain level, it generated a number of additional chains when compared to the native chains. Furthermore, the chain paths in the structures predicted by CryoREAD were markedly divergent from those in the native structures. The comprehensive benchmark test results are detailed in the Results section.

To address the challenges outlined above, we have developed a novel deep learningbased approach termed “DeepCryoRNA” for fully automated reconstruction of RNA 3D structures from protein-free cryo-EM maps with resolutions better than 6 Å. DeepCryoRNA utilizes a trained U-Net neural network [26, 27, 30] to predict 18 types of RNA atoms, including 12 backbone atoms and 6 base atoms. It then performs dedicated post-processing to connect these atoms into nucleotides, link nucleotides into short chains, extend short chains into long chains, and assign RNA sequences through modified global sequence alignments. Following the post-processing steps, energy-minimized all-atom RNA 3D structures are generated. In contrast to CryoREAD [25], our DeepCryoRNA method reported here demonstrates a broader range of atom type predictions using the U-Net neural network, thereby enhancing the nucleotide assembly process from the predicted atoms. Unlike CryoREAD, which identifies all four base types, the current DeepCryoRNA simplifies this to only two base types (purines and pyrimidines). This simplification is attributed to the inherent challenge encountered by neural networks in effectively discriminating between adenine and guanine, as well as between cytosine and uracil from cryo-EM maps of medium to low resolutions. Notably, DeepCryoRNA assigns RNA sequences through global sequence alignments, while CryoREAD relies on fragment sequence alignments. Furthermore, given the scarcity of pure RNA structures solved by cryo-EM, we opt to utilize RNA structures extracted from RNA-protein complexes to train the atom U-Net neural network. When benchmarked against the 51 test RNAs with cryo-EM map resolutions better than 6.0 Å, DeepCryoRNA demonstrates clear significant improvements over existing methods [24, 25].

## 2 Results

### 2.1 An overview of DeepCryoRNA

DeepCryoRNA aims to reconstruct RNA 3D structures from provided cryo-EM maps and corresponding RNA sequences. This procedure entails data preprocessing, atom prediction using a trained neural network, and the subsequent postprocessing of atoms to form nucleotides, short chains, and long chains. Ultimately, the native RNA sequence is precisely assigned to the postprocessed chains through global sequence alignments. It is noteworthy that to augment the database for the training of atom neural network, we included RNA molecules within the RNA-protein complex structures. A total of 131 RNA structures were extracted from the RNA-protein complexes and used for training. The detailed information for the training RNAs and cryo-EM maps is documented in Supplementary Table 1 for reference. Fig. 1 provides an overview of the primary steps in DeepCryoRNA, which will be briefly introduced here and elaborated in the Methods section.

**Fig. 1.**
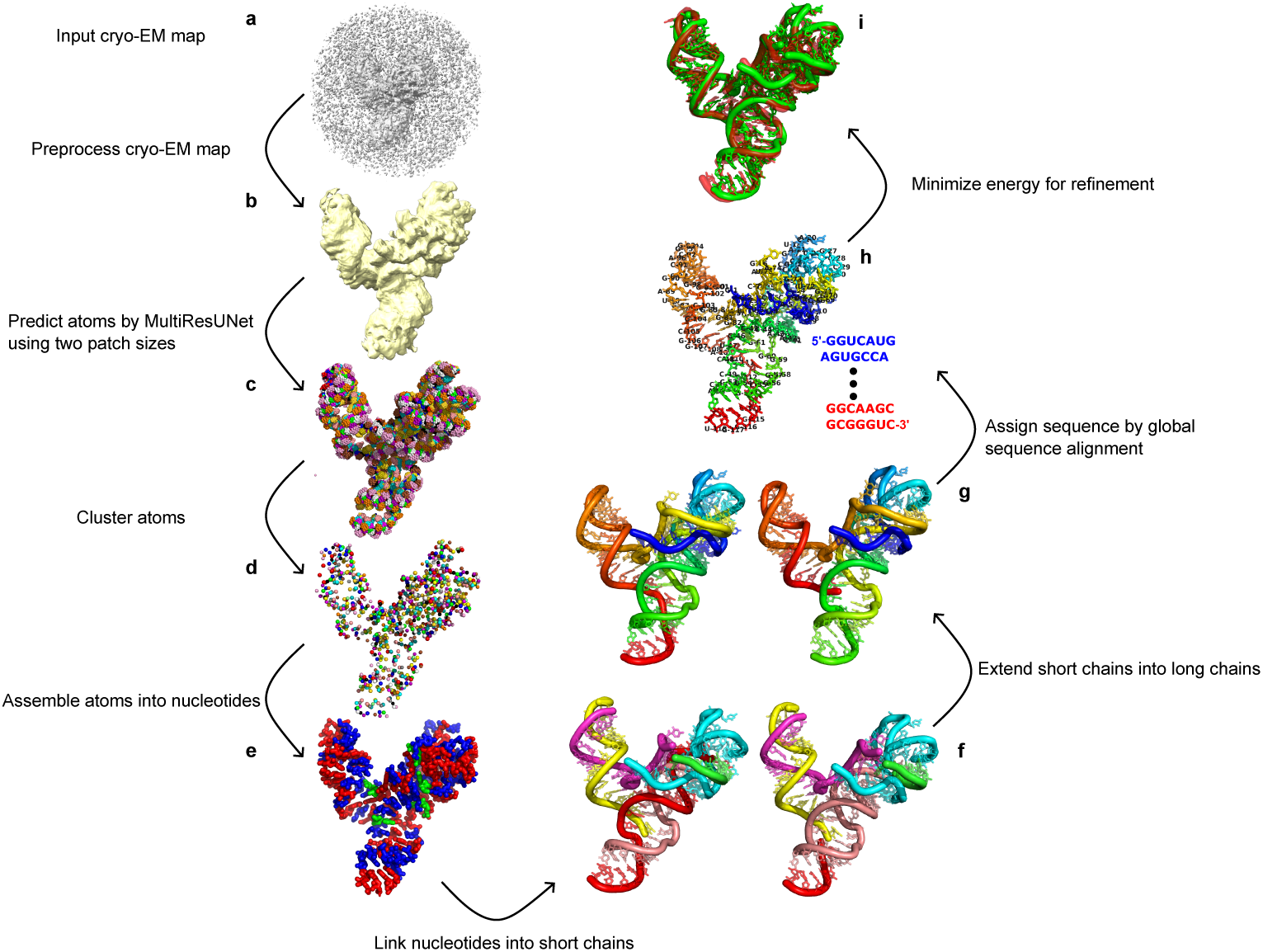
Flowchart of DeepCryoRNA. **a,** The raw cryo-EM map downloaded from the EMDB. **b,** Preprocessing of the cryo-EM map, involving resampling and normalization. **c,** Atom prediction via our trained MultiResUNet neural network, utilizing two patch sizes, 128 and 64. Predicted RNA atoms are represented by spheres, with different colors indicating distinct atom classes (18 in total). **d,** Atom clustering to eliminate redundant predicted atoms, with different atom types indicated by color. **e,** Nucleotide assembly from the clustered atoms, yielding three nucleotide types: A (red), C (blue), and X (green). **f,** Building short RNA chains by linking bonded nucleotides (until no map-derived nucleotides can be added to the chain). This example illustrates the formation of two distinct sets of short chains, with each individual short chain color-coded. **g,** Extending nearby short chains to create long RNA chains. Two sets of long chains are formed in this example, and they are colorfully displayed with a rainbow scheme, featuring a blue 5’-end and a red 3’-end. **h,** Assignment of native sequences through global sequence alignments. Accurate nucleotide indexing and chain information is assigned based on the best sequence alignment. **i,** Energy minimization to refine the complete all-atom RNA 3D structure using the QRNAS software. Both the native (red) and reconstructed (green) structures are shown.

#### 2.1.1 Preprocessing data

During both the neural network training and prediction phases, it is crucial to perform preprocessing on the raw cryo-EM maps obtained from the EMDB [28]. This preprocessing entails two key steps: map resampling to a fixed voxel size of 0.5 Å by the software ChimeraX [31] and map data normalization to ensure a consistent data range. In the neural network training phase, it is also necessary to preprocess the RNA structures within the training dataset. This preprocessing is aimed at generating atom class maps that correspond to the cryo-EM maps. The neural network is designed to predict 19 distinct atom classes, including one for background atoms and 18 for RNA atoms. More specifically, these RNA atom classes consist of 12 RNA backbone atoms (P, OP1, OP2, C5’, O5’, C4’, O4’, C3’, O3’, C2’, O2’, C1’) and 6 RNA base atoms (N9, C4, C8 for purines, and N1, C2, C6 for pyrimidines).

#### 2.1.2 Atom prediction and clustering

The trained MultiResUNet neural network [30], a variant of the classical U-Net architecture [26, 27], is applied to predict atom classes from a preprocessed cryo-EM map. Specifically, a complete cryo-EM map is divided into multiple patches, and each cryo-EM map patch is fed into the neural network to predict atom classes. Subsequently, the predicted atom class map patches are interconnected to reconstruct the overall atom class map. Two patch sizes, namely 64^3^ and 128^3^, serve as input dimensions for the neural network tasked with atom class prediction. For brevity, we refer to them as “patch size 64” and “patch size 128” in this context. Such two patches result in two sets of predicted atoms denoted as “atom-ps64” and “atom-ps128”, respectively. During the neural network training phase, a single patch size of 64 was utilized, and different patches of size 64 were randomly cropped from the complete cryo-EM maps in the training dataset. During the prediction stage, we diversify the map input information by employing two different patch sizes. The neural network training and prediction were conducted in the open-source machine learning framework TensorFlow [32]. Furthermore, the predicted atoms undergo clustering to eliminate redundancy of the predictions from nearby voxels, as shown in Fig. 1c and d.

#### 2.1.3 Nucleotide assembly from atoms

Nucleotides are constructed based on the clustered atoms, considering their atom classes and pairwise atomic distances. Nucleotide types are identified by examining the base atom classes and their quantities. Notably, our approach distinguishes between purines and pyrimidines while not discerning the four RNA nucleotide types (A, C, G, U). As a result, predicted nucleotides are designated as either Adenine (A), Cytosine (C), or an unknown type (X). Two sets of nucleotides, denoted as “nt-ps64” and “nt-ps128”, are derived from the respective sets of atoms “atom-ps64” and “atom-ps128”.

#### 2.1.4 Formation of short and long chains from nucleotides

Short chains are created by linking neighboring nucleotides, and it’s important to note that a single nucleotide can have multiple neighboring nucleotides, resulting in the generation of multiple sets of short chains from a set of nucleotides. Within these sets of short chains, long chains are formed by connecting neighboring short chains. Just like a nucleotide, a single short chain can have multiple neighboring short chains, leading to the creation of multiple sets of long chains from a given set of short chains. Each set of long chains constitutes a complete chain, and it is feasible to derive a multitude of complete chains, each corresponding to distinct connection pathways among the nucleotides and short chains. Evidently, two sets of short and long chains are acquired for the two distinct patch sizes.

#### 2.1.5 Native sequence assignment by global sequence alignment

Global sequence alignments are employed to align the complete long chains with the provided native RNA chains. We use a modified Gotoh algorithm [33] for these alignments, with a key feature being the utilization of distinct gap opening and extension costs at different positions. The intent of this design is to apply a more substantial penalty to gap insertions within the spatially continuous short chains, thereby preserving the short chains’ structural integrity. From the top 10 best alignment results, we assign native chain information to the corresponding complete long chains, ultimately yielding 10 all-atom RNA structures for each of the two sets of nucleotides “nt-ps64” and “nt-ps128”. Furthermore, we identify the overall top 10 predicted structures by evaluating the alignment scores for both patch sizes. Finally, structural refinement is carried out through the software QRNAS [34] to rectify broken bonds and steric clashes between atoms in the predicted RNA structures. In summary, the approach yields up to 10 predicted structures for each category denoted as “RNA-ps64”, “RNA-ps128”, and “RNA-psBoth”. DeepCryoRNA encompasses three distinct versions: DeepCry-oRNA ps64, DeepCryoRNA ps128, and DeepCryoRNA psBoth, each corresponding to methods utilizing a single patch size of 64 and 128, and both patch sizes of 64 and 128, respectively. In the practical implementation, users need not specify the methods, as the predicted structures from all three methods are provided if available.

### 2.2 Comparative performance analysis

We conducted comparisons between our method, DeepCryoRNA psBoth, and two other deep learning-based methods, DeepTracer [24] and CryoREAD [25], to demonstrate each method’s capability in reconstructing RNA 3D structures from cryo-EM maps. Furthermore, we compared DeepCryoRNA psBoth with DeepCryoRNA ps64 and DeepCryoRNA ps128 to investigate the effect of patch size on prediction results. The benchmark test dataset consists of 51 pure RNAs with cryo-EM maps of resolutions better than 6.0 Å. The detailed information for the 51 test RNAs and cryo-EM maps is shown in Supplementary Table 2. We employed the DeepTracer and CryoREAD servers to perform RNA structure predictions using cryo-EM maps. The comparison between all the methods were based on the same input data, including the raw cryo-EM map obtained from EMDB [28], the respective RNA sequence in FASTA format, the contour level and cryo-EM map resolution sourced from the EMDB. The results of the comparison between these methods are presented in Fig. 2 and detailed in Supplementary Table 3. The performance comparison is based on the RMSD between the native RNA 3D structures and the structures predicted by each method, as illustrated in Fig. 2. The RMSD calculations in this study were based on all the heavy atoms and conducted using the “align” command in the software PyMOL [35]. Both DeepTracer and CryoREAD yielded a single predicted structure for each RNA. In contrast, each of our three approaches has the capacity to generate up to 10 predicted structures per RNA. For the purpose of comparison, we selected the structure of the highest global sequence alignment score obtained in the native sequence assignment process for each of our approaches.

**Fig. 2.**
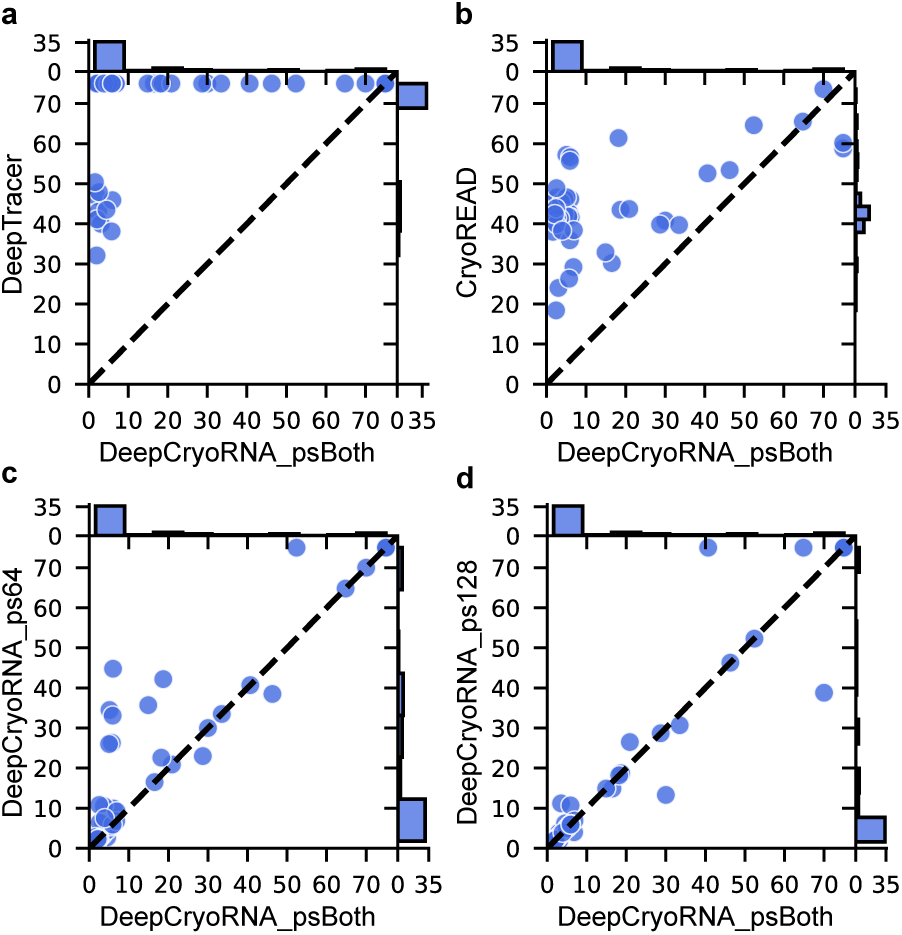
Performance comparison of RMSD between DeepCryoRNA psBoth and four alternative methods: **(a)** DeepTracer, **(b)** CryoREAD, **(c)** DeepCryoRNA ps64, and **(d)** Deep-CryoRNA ps128. Methods DeepCryoRNA psBoth, DeepCryoRNA ps64, and DeepCryoRNA ps128 are our approaches employing two patch sizes of 64 and 128, a single patch size of 64, and 128, respectively. In each subfigure, the x-axis and y-axis of the scatterplot represent the RMSD values (in Å) between the native RNA 3D structures and the corresponding predicted structures obtained from the two methods, as denoted by the x- and y-labels. The blue solid circles indicate the 51 RNAs in the test dataset. The RMSD values span from 0 to 75 Å, where 75 Å signifies cases in which the methods fail to predict structures. Histogram plots depicting the RMSD distributions are presented along the top and right sides in each subfigure.

We compared the performance of DeepCryoRNA psBoth with the existing method, DeepTracer [24], as depicted in Fig. 2a. We found that for 41 out of the 51 instances, DeepTracer failed to predict the structure either because only phosphorus atom positions were predicted or only a fraction of nucleotides in the native RNA structures were included, or the server itself malfunctioned. In contrast, DeepCryoRNA psBoth achieved low RMSD values for most RNA structures, with 36 RNAs ranging from 1.6 to 6.9 Å, 5 RNAs from 14.9 to 20.9 Å, and 8 RNAs from 28.7 to 70.0 Å. The average and median RMSDs for DeepCryoRNA psBoth are 12.1 and 5.0 Å, respectively. As shown in Fig. 2b, CryoREAD could successfully provide structure prediction for all the 51 test RNAs. However, the resulting RMSDs are notably high, with values ranging from 18.4 to 73.6 Å. The average and median RMSDs are 44.3 and 42.4 Å, respectively. These result indicate a substantial improvement of the DeepCryoRNA psBoth approach.

In addition to conducting comparisons with the existing methods, we also evaluated the performance of DeepCryoRNA psBoth and its two variants: DeepCryoRNA ps64 and DeepCryoRNA ps128, which employ a single patch of size 64 and 128, respectively. We found that DeepCryoRNA psBoth outperformed DeepCryoRNA ps128, as evident in Fig. 2d, while exhibiting markedly better performance than DeepCryoRNA ps64, as depicted in Fig. 2c. This result demonstrates that the utilization of two patch sizes yields better results in comparison to a single patch size, as it enhances the probability of identifying the correct chain paths. The rationale behind the better performance of DeepCryoRNA ps128 than DeepCryoRNA ps64 lies in the fact that a larger patch size (128) allows the neural network to encompass a more extensive portion of the cryo-EM map, facilitating the prediction of atom information, especially in regions proximate to the patch boundaries. Additionally, a larger patch size reduces the overall number of patches, thereby mitigating the impact of patch boundary effects on the atom prediction process. Furthermore, our evaluation, as depicted in Figs. 2c and d, reveals instances in which DeepCryoRNA psBoth underperforms in comparison to either DeepCryoRNA ps64 or DeepCryoRNA ps128. This underscores the necessity of considering the results from DeepCryoRNA ps64 and DeepCryoRNA ps128 in practical applications.

Fig. 3 shows eight examples of RNA structures predicted by DeepCryoRNA, Deep-Tracer, and CryoREAD, compared with the corresponding native structures. The cryo-EM maps for the three RNAs (PDB 7EZ2 [36], 6UES [37], 7XD3 [38]) in Figs. 3a-c exhibit resolutions of 3.05, 3.7, and 4.05 Å, respectively. Our approach, DeepCryoRNA, produced structures that exhibit high similarity to the corresponding native structures, featuring low RMSDs of 2.0, 2.4, and 2.5 Å, respectively. In contrast, the structures generated by DeepTracer and CryoREAD display larger RMSDs to the native structures. Specifically, DeepTracer resulted in numerous fragmented chains for RNA 7EZ2 in the third column of Fig. 3a, and in the case of RNAs 6UES and 7XD3, as depicted in Figs. 3b and c, it only generated a limited number of nucleotides. In contrast to DeepTracer, CryoREAD predicted a larger number of nucleotides, but these nucleotides were distributed across a considerably larger number of chains than observed in the native structures. Consequently, the predicted nucleotides cannot align with their counterparts in the native structures. This mismatch implies that two nucleotides occupying nearly identical positions have different nucleotide indices and chain information.

**Fig. 3.**
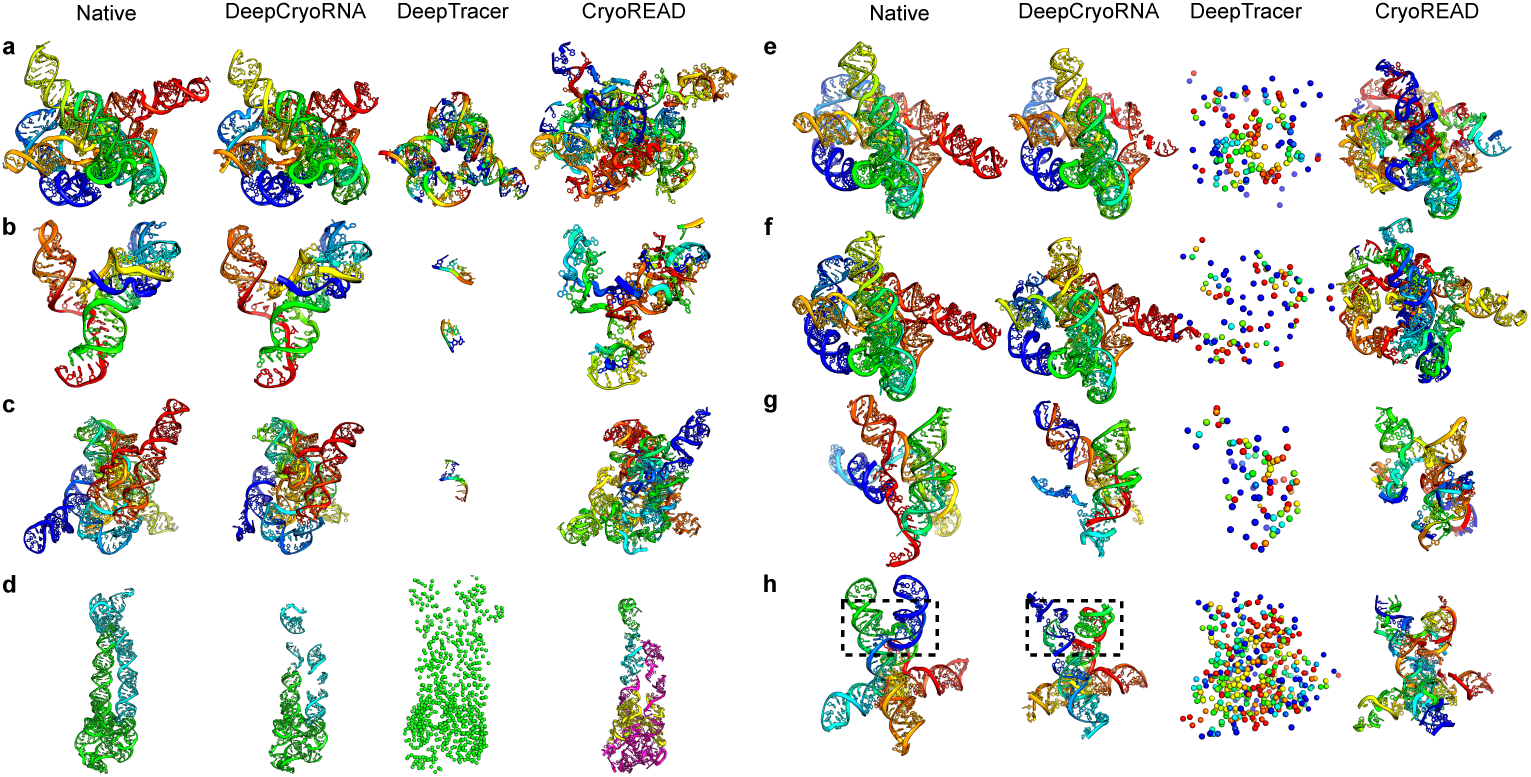
Examples of the RNA structures predicted by DeepCryoRNA, DeepTracer, and CryoREAD. Each panel consists of four columns, representing the native RNA structure and structures predicted by DeepCryoRNA, DeepTracer, and CryoREAD, respectively. Each chain in the structures is color-coded in a rainbow spectrum, with the 5’-end represented in blue and the 3’-end in red. In panel d, green and cyan colors are used to distinguish chains A and B, respectively. The visual representation employs a cartoon style, except for the structures predicted by DeepTracer in panels d-h, which are shown as spheres to indicate that only phosphorus atoms are predicted in those structures. This color-coding scheme is utilized to enhance the visual discernment of structural comparisons between the predicted and native RNA structures. **a,** RNA 7EZ2 with map resolution of 3.05 Å. RMSD: 2.0 Å (DeepCryoRNA), 46.9 Å (DeepTracer), 45.7 Å (CryoREAD). **b,** RNA 6UES with map resolution of 3.7 Å. RMSD: 2.4 Å (DeepCryoRNA), NaN (DeepTracer), 18.4 Å (CryoREAD). **c,** RNA 7XD3 with map resolution of 4.05 Å. RMSD: 2.5 Å (DeepCryoRNA), NaN (DeepTracer), 48.9 Å (CryoREAD). **d,** RNA 8SA5 with map resolution of 3.5 Å. RMSD: 18.7 Å (DeepCryoRNA), NaN (DeepTracer), 43.5 Å (CryoREAD). **e,** RNA 7XSM with map resolution of 4.01 Å. RMSD: 13.3 Å (DeepCryoRNA), NaN (DeepTracer), 40.8 Å (CryoREAD). **f,** RNA 7YC8 with map resolution of 4.14 Å. RMSD: 20.9 Å (DeepCryoRNA), NaN (DeepTracer), 43.7 Å (CryoREAD). **g,** RNA 7SAM with map resolution of 4.3 Å. RMSD: 16.5 Å (DeepCryoRNA), NaN (DeepTracer), 30.2 Å (CryoREAD). **h,** RNA 8BTZ with map resolution of 5.39 Å. RMSD: 28.7 Å (DeepCryoRNA), NaN (DeepTracer), 39.8 Å (CryoREAD). Note: “NaN” indicates an unsuccessful prediction.

As depicted in Fig. 2 and Supplementary Table 3, DeepCryoRNA achieved RMSDs equal to or less than 6.0 Å for 35 out of the 51 test RNAs, demonstrating its effectiveness. Upon examining cases with larger RMSDs, we identified that the challenges primarily arise from limitations in the heterogeneous, low-resolution local regions of cryo-EM maps. Notably, omitting such problematic regions enables the attainment of smaller RMSDs. To gain deeper insights into these challenges, as shown in Figs. 3d-h, we provide illustrative examples of five cases for which DeepCryoRNA provides relatively poor predictions.

In the case of RNA 8SA5 [39] depicted in Fig. 3d, the RNA adopts a dimeric structure with chains A and B distinguished by green and cyan colors, respectively. The cryo-EM map corresponding to this RNA exhibits a resolution of 3.5 Å. Our approach yielded a relatively large RMSD of 18.7 Å compared with the complete dimeric structure. Notably, when examining chain A, the predicted structure closely mirrors the native one, showcasing an RMSD of 2.6 Å. The elevated RMSD for the entire dimer structure primarily arises from chain B. An examination of Fig. 3d (second and fourth columns) reveals that DeepCryoRNA and CryoREAD predicted only a limited segment of nucleotides in chain B, which is colored in cyan in the native structure. This observation implies that the cryo-EM map for RNA 8SA5 possesses a heterogeneous resolution, with a notably lower resolution in the region corresponding to chain B. DeepTracer only produced a cluster of phosphorus atoms, as depicted in the third column of Fig. 3d. The chain trajectories predicted by CryoREAD exhibit notable deviations from the native paths shown in the fourth column of Fig. 3d, leading to a substantial RMSD of 43.5 Å.

In the case of RNA 7XSM [40] with a cryo-EM map resolution of 4.05 Å, as illustrated in Fig. 3e, our approach, DeepCryoRNA, achieved an RMSD of 13.3 Å. Upon comparing our predicted model with the native structure, we identified that the elevated RMSD primarily stems from nucleotides 326-388 located in the 3’-end of the RNA. Specifically, for nucleotides 1-325, the RMSD between our prediction and the native structure is 4.2 Å, whereas for nucleotides 326-388, the RMSD increases to 29.4 Å due to an incorrect chain path caused by a large number of missing nucleotides in this region. Consequently, for this RNA, our approach accurately predicted the structure for the majority of the sequence, with the exception of the dozens of trailing nucleotides. As in the previous situation, DeepTracer continued to yield only phosphorus atoms, while CryoREAD produced numerous additional chains and nucleotides that did not align with the native structure. In the case of RNA 7YC8 [41] with a cryo-EM map resolution of 4.14 Å, depicted in Fig. 3f, our DeepCryoRNA approach yielded a substantial RMSD of 20.9 Å. Much like the previous RNA, 7XSM, this elevated RMSD results from an incorrect chain path in part of our predicted structure. Notably, for approximately 200 out of the 333 predicted nucleotides, the RMSD is a more modest 3.1 Å, signifying that, while the overall predicted structure exhibits a considerable RMSD, a portion of the predicted structure aligns accurately with the native one. Despite partial inaccuracies, this pre-dicted structure can assist users in manual corrections, especially when complemented by secondary structures. Similar to the preceding example, in this case, DeepTracer generated only phosphorus atoms, and CryoREAD produces a significant RMSD of 43.7 Å.

In the case of RNA 7SAM [42], as depicted in Fig. 3g, our approach yielded a large RMSD of 16.5 Å, primarily due to the heterogeneous resolution in the cryo-EM map. While the overall resolution of the cryo-EM map for this RNA is 4.3 Å, the map resolution for the 5’-end, 3’-end, and the central region of the RNA is considerably lower, rendering both DeepCryoRNA and CryoREAD unable to predict the nucleotides in those regions. Furthermore, DeepTracer continued to predict only phosphorus atoms for this RNA.

In the case of RNA 8BTZ [43], as depicted in Fig. 3h, our method resulted in a large RMSD of 28.7 Å. This RNA is characterized by four-way junctions, and all the junction loop lengths are zero, as delineated within the dashed rectangle in the first column of Fig. 3h. However, our DeepCryoRNA approach predicted it as two separate helices, as indicated by the dashed rectangle in the second column of Fig. 3h. Notably, discerning a four-way junction without junction loops from a cryo-EM map with a resolution of 5.39 Å can be challenging. As before, DeepTracer produced solely phosphorus atoms, while CryoREAD yielded a large RMSD of 39.8 Å.

In summary, our approach DeepCryoRNA outperformed DeepTracer and CryoREAD in reconstructing RNA 3D structures from RNA-only cryo-EM maps. DeepTracer frequently failed to generate valid structures, whereas CryoREAD, while reconstructing nucleotides, deviated significantly from native chain paths and introduced extra chains. In contrast, DeepCryoRNA consistently and accurately reconstructed RNA structures from cryo-EM maps of high resolutions, with performance variation observed for cryo-EM maps of low resolutions. The variability in performance in some cases can be attributed to heterogeneous map resolutions. Even in cases where predictions were less accurate, certain structural components can still be correct, offering valuable insights for human intervention.

### 2.3 Performance analysis of DeepCryoRNA

In accordance with the data presented in Fig. 2, our DeepCryoRNA psBoth approach demonstrates structure reconstruction with RMSDs ranging from 1.6 to 70.0 Å for 49 test RNAs and achieves low RMSDs for the majority of the RNAs. This result prompts an investigation into the underlying factors contributing to the performance. We have conducted an analysis of the relationship between RMSDs and five potential determinants affecting prediction accuracy, as illustrated in Fig. 4. The distributions of RMSDs and the respective influential factors are also depicted in each subfigure.

1. Fig. 4a illustrates the relationship between the resolution of cryo-EM maps and RMSD for the 51 test RNAs. Notably, a significant increase in RMSDs is observed when the cryo-EM map resolution is low (beyond 4.5 Å). For such low-resolution EM maps, DeepCryoRNA is capable of achieving reliable structure models for map resolutions greater than 4.5 Å, although the performance is not as consistent as it is for higher cryo-EM map resolutions. It is noteworthy that in cases where cryo-EM maps with resolutions better than 4.5 Å result in large RMSDs, the majority of a sequence continue to exhibit low RMSDs, although the heterogeneity in cryo-EM maps may lead to incorrect chain paths in specific sections for certain cases. These results suggest that our method enables reliable reconstruction of RNA structures when provided with cryo-EM maps of resolutions better than 4.5 Å, while predictions show variable performance for cryo-EM maps of lower resolutions.
2. Fig. 4b shows the relationship between RNA length and RMSD across the 51 test RNAs. It is evident that large RMSDs could occur for RNAs of various sizes, indicating that the performance is not sensitive to RNA length. Instead, the performance is more influenced by the cryo-EM map resolution, as shown by Fig. 4a and b.
3. In Fig. 4c, we explore the relationship between the probability of successfully rebuilding nucleotides and the RMSD for the 51 test RNAs. This probability is computed as the ratio of the number of rebuilt nucleotides to the total number of nucleotides in the native RNA structure, without considering the correctness of nucleotide types. Generally, a higher probability of nucleotide rebuilding corresponds to lower RMSDs in our approach. When this probability exceeds 85%, we consistently achieve low RMSDs; otherwise, the RMSDs are more variable. Even in instances of less successful predictions, our approach maintains a nucleotide rebuilt probability exceeding 60%, thereby providing valuable structural information to aid users in further refining RNA structures.
4. Fig. 4d explores the relationship between the accuracy of predicting nucleotide types and RMSD for the 51 test RNAs. Notably, a prediction accuracy exceeding 80% consistently results in lower RMSDs, while accuracy below 80% is associated with varying RMSDs. High prediction accuracy of nucleotide types facilitates precise sequence alignments and the identification of correct chain paths.
5. In Fig. 4e, we examine the relationship between the number of the predicted short chains and RMSD for the 51 test RNAs. Generally, the presence of more short chains in predictions, indicating the division of native RNA structures into numerous segments, complicates the task of threading these short chains into the correct complete long chain of the native RNA. The presence of additional short chains often coincides with more missing nucleotides between these segments, rendering the accurate connection of short chains and sequence assignment by alignment challenging.

**Fig. 4.**
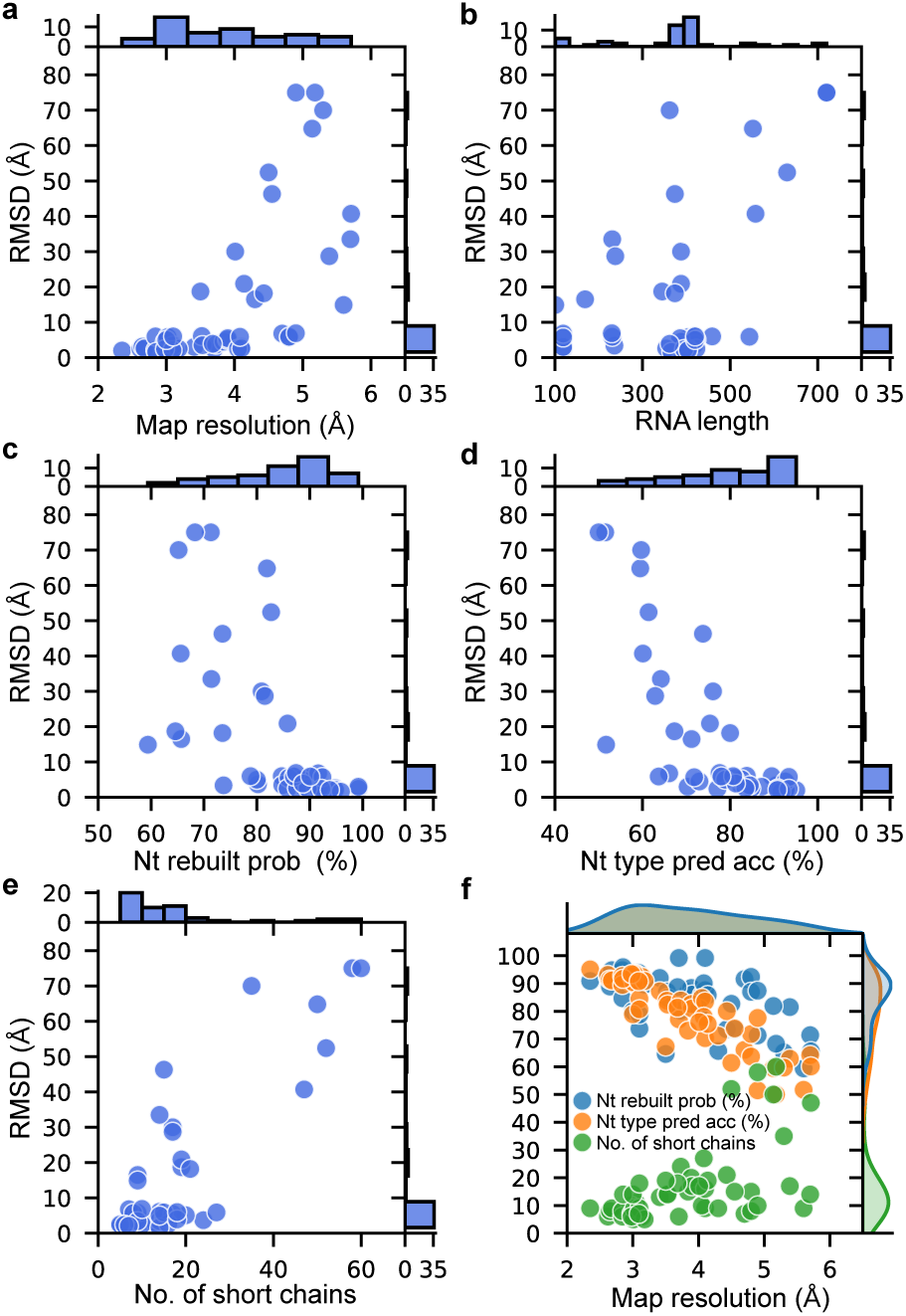
Factors affecting DeepCryoRNA performance. In each subfigure, the y-axis of the scatterplot represents RMSD achieved by the DeepCryoRNA psBoth method. Histogram plots depicting the influencing factor and RMSD distributions are presented along the top and right sides in each subfigure, respectively. **a,** Cryo-EM map resolution vs. RMSD. **b,** Native RNA length vs. RMSD. **c,** Nucleotide rebuilt probability vs. RMSD. **d,** Nucleotide type prediction accuracy vs. RMSD. **e,** Number of short chains vs. RMSD.

Furthermore, it is important to note that the factors above are not mutually exclusive, and the latter three factors exhibit a strong correlation with cryo-EM map resolutions, as illustrated in Fig. 4f. The results indicate that cryo-EM maps of higher resolutions are associated with an increased likelihood of successful nucleotide rebuilding, improved accuracy in predicting nucleotide types, and a reduction in the number of predicted short chains.

### 2.4 Effective modified global sequence alignment

To assign the correct native chain information to the predicted complete long chains, we conducted a global sequence alignment between the query sequences derived from the predicted chains and the reference native chains. This alignment was performed using a customized Gotoh algorithm, which penalizes more severely gaps inserted within spatially continuous short chains. Additionally, we used the same alignment algorithm but applying uniform gap penalties regardless of gap positions. The results were then compared with those obtained using our DeepCryoRNA approach, which takes gap positions into account; see Fig. 5, where RMSD1 and RMSD2 were calculated for the structures predicted with and without considering gap position-dependent penalties, respectively. Fig. 5 shows that RMSD1 is generally lower than RMSD2, particularly in regions with low RMSD1 values. The modified global sequence alignment approach facilitates the preservation of intact short chains, particularly those comprising a number of nucleotides with inaccurately predicted nucleotide types.

**Fig. 5.**
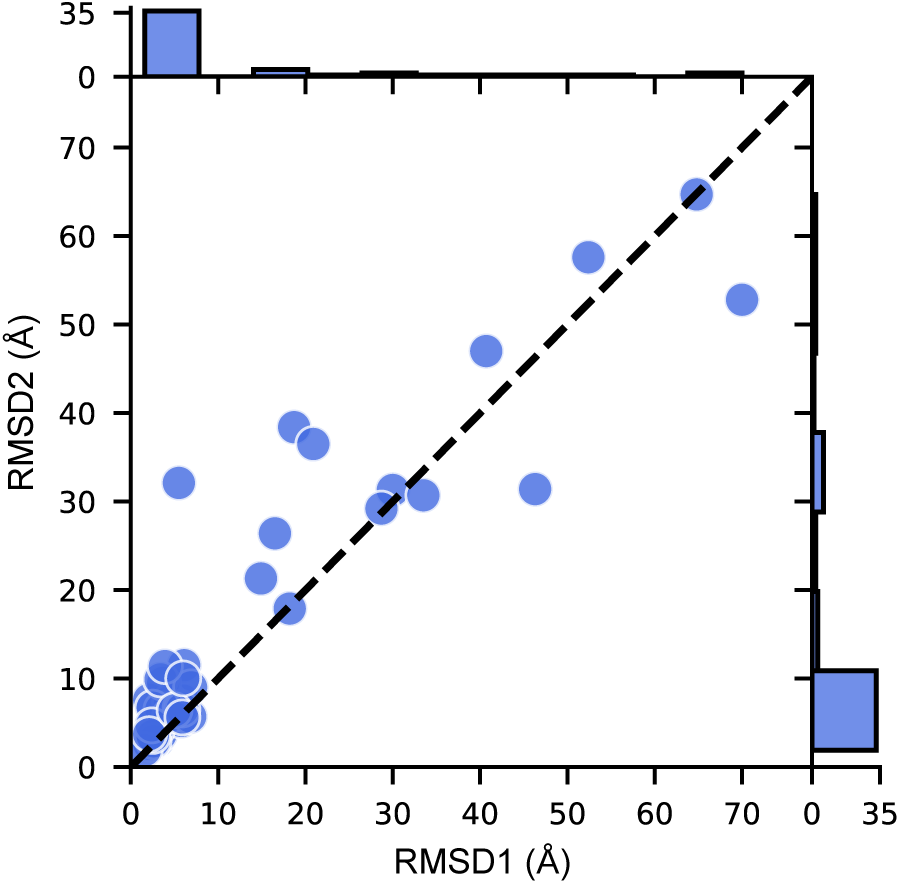
**Comparison of RMSDs achieved by two alignment methods.**RMSD1 represents the RMSD calculated using DeepCryoRNA’s alignment method, while RMSD2 represents the RMSD calculated using the conventional alignment method. The histogram plots show the RMSD distributions.

### 2.5 Runtime

We used a GPU (NVIDIA GeForce GTX 1080 Ti) and 10 CPUs (AMD Ryzen Threadripper, 3.4 GHz) to conduct our approach DeepCryoRNA for the 51 test RNAs. The relationship between RNA length and runtime is shown in Fig. 6. The runtimes range from 4 minutes to 3.5 hours.

**Fig. 6.**
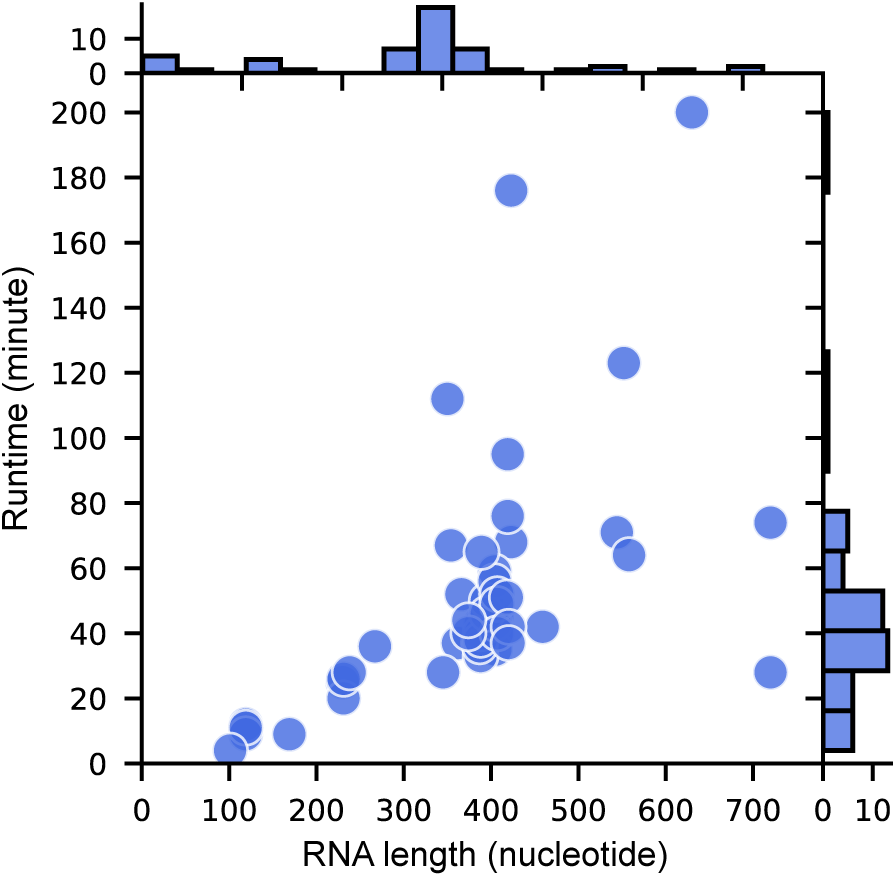
The relationship between RNA length and runtime for the 51 test RNAs. Their respective distributions are also shown in the histogram plots.

## 3 Discussion

We have developed DeepCryoRNA, an advanced deep learning-based methodology designed for the *de novo* reconstruction of RNA 3D structures from RNA-only cryo-EM maps and sequences. The reconstruction process encompasses atom prediction through a trained neural network, followed by meticulous post-processing. During post-processing, predicted atoms are organized into nucleotides, which are subsequently linked to form short chains, and these short chains are further extended into long chains. Finally, native sequence information is assigned to the extended chains through global sequence alignments.

In the case of cryo-EM maps with resolutions finer than 4.5 Å, our method consistently yields near-native structures with low RMSDs. However, for maps with resolutions below 4.5 Å, the performance may exhibit variability, as depicted in Fig. 4a. In contrast to conventional conformational sampling-based methods, our approach is entirely automated, eliminating the requirement for secondary structure information, and excels in efficiency. When compared to the deep learning-based techniques DeepTracer [24] and CryoREAD [25], our method demonstrates substantially reduced RMSDs, as illustrated in Fig. 2a and b, primarily due to our capability to accurately determine the nucleotide identity, including the nucleotide index and the associated chain. The foundation of this impressive performance, when compared to the other two deep learning methods, lies in our meticulous postprocessing.

While DeepCryoRNA has demonstrated effectiveness overall, some limitations are evident in the current approach. Firstly, the method is exclusively applicable to pure RNAs and does not extend to DNAs or RNA-protein complexes. In contrast, Deep-Tracer can model proteins, DNAs, RNAs, and complexes, while CryoREAD can model DNAs and RNAs within protein-nucleic acid complexes. We anticipate that in future iterations, our approach will offer a unified solution for these diverse scenarios. Secondly, due to the inherent heterogeneity in cryo-EM maps, the predicted structures may lack certain loop conformations and, in some cases, helical structures. Users may leverage supplementary software, such as SimRNA [44, 45] and RNApps [46], to address missing loop elements and refine structures using tools like Phenix [12] or correlation-driven molecular dynamics. Incorporating these enhancements would further help streamline the final modeling process. As cryo-EM techniques continue to advance, an increasing number of pure RNA structures are being resolved using this technology. We anticipate that DeepCryoRNA will prove to be a powerful tool for reconstructing RNA structures in this evolving landscape.

## 4 Methods

### 4.1 Training dataset collection

In constructing our training dataset, we utilized RNA structures from RNA-protein complex structures by following a systematic approach. Initially, we retrieved RNA-protein complex structures determined through cryo-EM with a resolution better than 7 Å from the RCSB PDB [29] as of February 2022. We then extracted the RNA structures from these complexes, excluding those with a length shorter than 50 nucleotides and removing redundant RNA structures using an RMSD threshold of 5 Å. This process yielded a total of 131 distinct RNA structures. Additionally, for each of these 131 RNA-protein structures, we obtained the corresponding cryo-EM maps from the EMDB [28] and used the command “volume zone” in the software Chimerax [31] to crop the map region within 5 Å of any atoms in the RNA structures, thereby elimnating the map volume occupied by protein structures. Consequently, our training dataset includes 131 RNA structures, with sequence lengths ranging from 50 to 15,871 nucleotides, each associated with a cryo-EM map exhibiting resolutions between 2.0 and 7.0 Å. The information for the RNAs and cryo-EM maps in the training dataset can be found in Supplementary Table 1.

### 4.2 Test dataset collection

In our test dataset, we included RNA structures devoid of any protein or DNA, which were resolved using cryo-EM with a resolution surpassing 6 Å and had been deposited in the RCSB PDB [29] prior to August 2023. This dataset encompasses 51 such RNA structures for which we also acquired the corresponding cryo-EM maps from the EMDB [28]. As a result, our test dataset comprises 51 pure RNA structures, with sequence lengths spanning from 101 to 720 nucleotides, each accompanied by a cryo-EM map showcasing resolutions between 2.3 and 5.7 Å. The information for the RNAs and cryo-EM maps in the test dataset can be found in Supplementary Table 2.

### 4.3 Data preprocessing

#### 4.3.1 Cryo-EM map preprocessing

Prior to inputting a cryo-EM map into the neural network, it must undergo crucial preprocessing steps, encompassing voxel resampling and data normalization. A cryo-EM map comprises a cuboid of voxels, each encapsulating electron density information. Notably, distinct cryo-EM experiments yield maps with varying voxel sizes, leading to a lack of uniformity in the spatial dimensions they represent. To resolve this issue and enable the neural network to assess inter-voxel distances accurately, we employed the “volume resample spacing 0.5” command in the software ChimeraX [31], resizing the voxels to have a consistent side length of 0.5 Å. Simultaneously, ensuring data consistency among cryo-EM maps is imperative, as they may exhibit diverse data ranges. This normalization process comprises several steps. First, the map data is truncated by capping all voxels at the 95th percentile of the electron density values to mitigate the influence of outliers on the neural network’s training. Subsequently, each voxel is modified by subtracting a specified contour level, while voxels smaller than 0 are reset to zero to eliminate artifacts associated with low electron density originating from solvent and noise. The contour level is adjusted to half of the EMDB’s recommendation. Lastly, all voxels are normalized by dividing them by the maximum electron density value, resulting in preprocessed cryo-EM map data ranging from 0 to 1. These meticulous preprocessing steps are vital for ensuring the reliability and efficacy of subsequent neural network training and predictions.

#### 4.3.2 RNA structure preprocessing for training

To train the neural network, we require the ground-truth voxel indices of atoms within RNA structures that correspond to the voxels in the cryo-EM maps. The calculation for determining the voxel indices of an RNA atom involves the following steps: Firstly, the origin coordinates of the corresponding cryo-EM map are subtracted from the atom’s coordinates. Subsequently, the atom’s coordinates are divided by 0.5, which represents the voxel size in the preprocessed cryo-EM map, and then rounded up to the nearest integers as the indices of the voxel corresponding to the atom. This voxel, along with its six directly neighboring voxels, is assigned the same atom class as the corresponding atom. After preprocessing the 131 RNAs in the training dataset, we obtained 131 atom class maps, akin to the cryo-EM maps, where each voxel represents an atom class. Our model employs 19 atom classes, encompassing 18 RNA atom classes and one additional atom class for the background, as elaborated in the main text.

### 4.4 Deep neural network: construction, training, and prediction

#### 4.4.1 Neural network construction

We utilized the MultiResUNet neural network [30], which is a variation of the conventional U-Net architecture [26, 27], to transform the cryo-EM maps into atomic representations. The architecture of the MultiResUNet used in our model is shown n Fig. 7. Compared to U-Net’s symmetric encoder-decoder structure, MultiResUNet utilizes a multi-resolution design with “Res Paths” to transform encoder features and “MultiRes Blocks” to aggregate multi-scale context, allowing it to leverage both local and global information for enhanced performance on image segmentation. This more complex architecture differentiates MultiResUNet from the simpler skip connection design of U-Net, while keeping model parameters comparable. In Fig. 7a, the 3D MultiResUNet architecture consists of 5 levels with a MultiRes (MR) block in each encoder and decoder level, along with 4 Res Paths between the encoder and decoder stages. The number of filters starts at 32 in the first level and doubles after each max pooling to 64, 128, 256, and 512 in the encoder path. The decoder path reverses this by halving the number of filters after each upsampling using transposed convolution operations. Within each MultiRes block as shown in Fig. 7b, the number of filters gradually increases from 1/6 to 1/3 to 1/2 of the block’s total filters. Res Paths 1 through 4 in Fig. 7a employ 32, 64, 128, and 256 filters in each Res Path Block in Fig. 7c, correspondingly. In Res Paths 1 through 4, there are 4, 3, 2, and 1 Res Path Blocks, respectively. Following the operations within the U-Net segment, a convolutional layer activated by the softmax function is employed to generate the probabilities of the 19 atom classes. The neural network consists of a total of 18,683,016 parameters, with 18,658,496 of them being trainable, and the remaining 24,520 parameters being non-trainable. We employed the TensorFlow Python package [32] to construct, train, and execute predictions with the neural network.

**Fig. 7.**
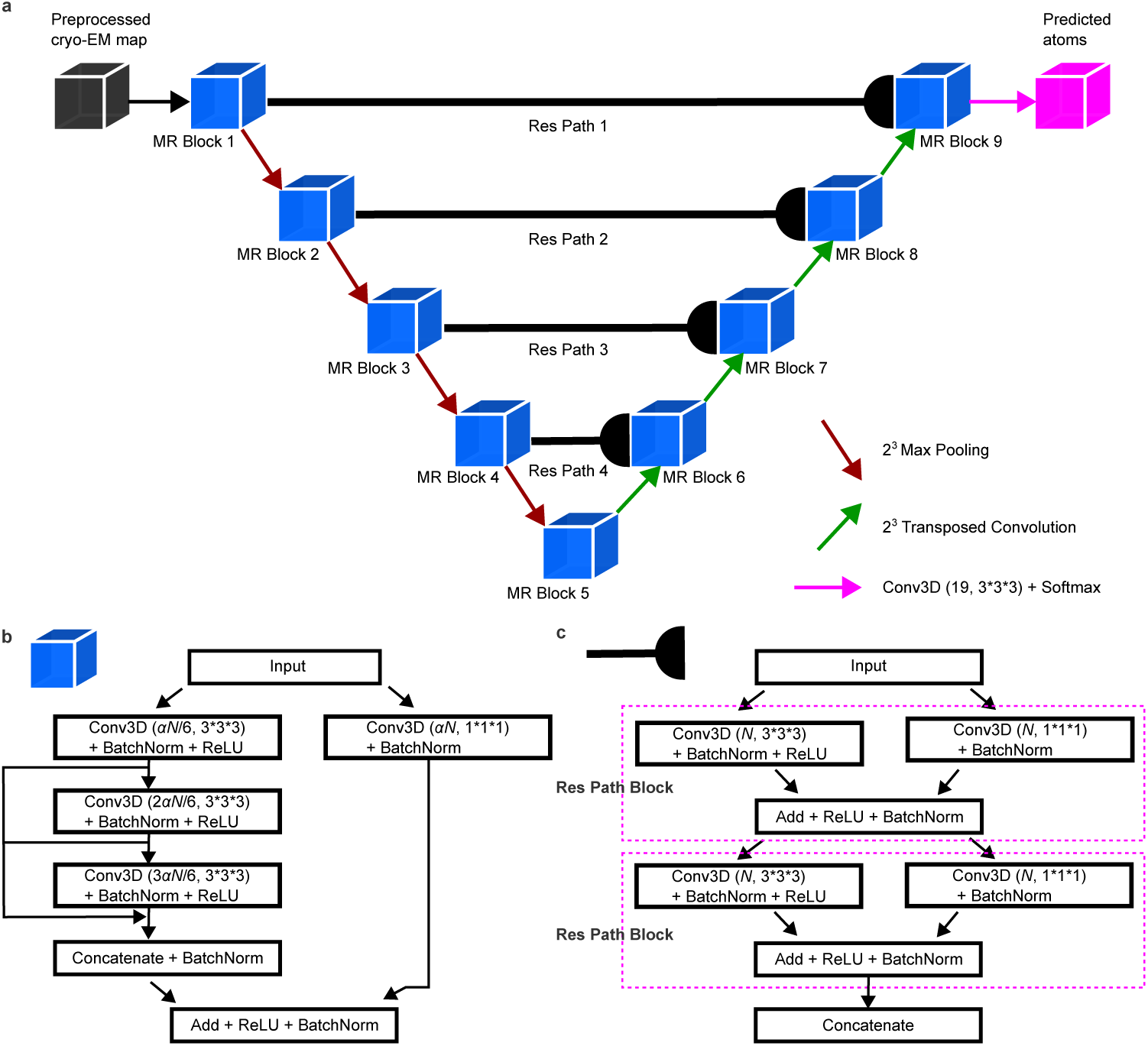
The MultiResUNet architecture used in our approach DeepCryoRNA. **a,** The overall structure of the neural network employs an encoder-decoder architecture, transforming a preprocessed cryo-EM map into a predicted atom class map. The encoder consists of five MultiRes (MR) blocks connected by 3D max pooling, while the decoder comprises five MR blocks connected by 3D transposed convolution. Four Res Paths link the encoder and decoder at different levels to extract and incorporate features at varying resolutions. **b,** Detailed view of a single MR block as presented in panel a. Here, Conv3D(*x*,*y*) represents a 3D convolution, with *x* denoting the number of filters and *y* specifying the filter size. The hyperparameter *α* is set to 1.67, in line with the original MultiResUNet model, and *N* represents the specified number of filters. BatchNorm denotes a batch normalization layer used to normalize inputs within a batch. ReLU, which stands for Rectified Linear Unit, serves as the activation function, outputting the input value if it’s positive and zero otherwise. The “Concatenate” operation fuses layers by expanding the feature dimension, while the “Add” operation performs element-wise addition of layers. **c,** A Res Path, as depicted in panel a, which can contain multiple Res Path Blocks enclosed within the dashed magenta rectangle. The annotations and labels used are consistent with those in panel b.

#### 4.4.2 Neural network training

In order to optimize the neural network, a mini-batch training methodology is utilized. This method allows us to efficiently process data and optimize model performance while mitigating overfitting. To generate diverse and informative mini-batches, we implemented a comprehensive data augmentation strategy. This process involves the following steps:

(1) Stochastic RNA selection: We begin by randomly selecting an RNA structure from the training dataset containing 131 diverse RNAs. The selection probability is proportional to the RNA structure’s volume, ensuring a representative training sample.
(2) Random patch cropping: For the selected RNA, we randomly crop a 64^3^ patch from the associated cryo-EM map, along with its corresponding RNA atom class map, ensuring that the cropped patch contains RNA atoms.
(3) Mixup augmentation: Next, we repeat the above process with another randomly selected RNA, creating a new patch. A crucial data augmentation technique, known as “mixup” is employed [47]. This involves blending the cryo-EM maps and atom class maps from two patches using the following equation 1, with *λ* ∈ [0, 1] representing the mixing coefficient randomly sampled from the Beta distribution Beta(2,2):

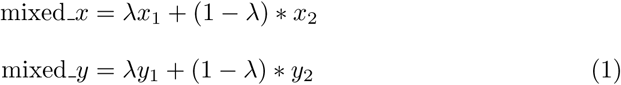

Here, *x*_1_ and *x*_2_ represent two randomly cropped cryo-EM map patches. The result, mixed *x*, is the newly created mixed cryo-EM map patch. Similarly, *y*_1_ and *y*_2_ correspond to the one-hot label encodings of the respective atom class map patches, while mixed *y* denotes the fresh label for the combined atom class map patch.

(4) Rotation augmentation: The new mixed patch is further augmented by randomly rotating it by 0, 90, 180, or 270 degrees along each axis.

These augmented 12 patches are then utilized to construct a mini-batch. During a single training epoch, we generated a total of 2730 such diverse mini-batches, with an average of approximately 250 crops per RNA.

To optimize training, we employed an atom-class-weighted Focal Loss [48] as the loss function. This involves setting the background atom class weight to 0.01, while assigning a weight of 1.0 to the 18 RNA atom classes. This weighting scheme prevents an undue emphasis on the dominant background atoms.

The neural network underwent 400 epochs of training, spanning approximately two weeks. The training process was conducted on three NVIDIA GeForce GTX 1080 Ti GPUs, each equipped with 11 GB of memory. We utilized the Adam optimizer [49] with an initial learning rate of 0.001, and the learning rate was halved if the training loss failed to decrease over 20 consecutive epochs, ultimately reaching a minimum learning rate of 0.0001.

#### 4.4.3 Neural network prediction

When applying our trained neural network to transform an RNA cryo-EM map into atomic classifications, a multi-step process is employed. Initially, the complete cryo-EM map is divided into smaller, adjacent patches. Atom class predictions are then generated for each of these individual map patches. Subsequently, these predicted atom class map patches are integrated to reconstruct the complete atom class map.

We employ two distinct patch sizes, denoted as *ps* = 64 and *ps* = 128, to partition the cryo-EM map into two separate sets of patches for prediction. It is worth noting that, in contrast to the random cropping employed during the training phase, our approach during the prediction stage involves cropping the cryo-EM map into adjacent and overlapped patches. This partitioning of the map introduces a potential challenge, as predictions in regions near the boundary of the patches might suffer from a lack of relevant information due to neighboring voxels being discontinuous. To mitigate this problem, we introduce a central core patch of dimensions (*ps* − 32)^3^ within each patch, following the approach described in the reference [15]. Although the dimensions of each patch remained *ps*^3^, we exclusively consider predictions from the inner core patch. As a result, during cryo-EM map patch generation, we incorporate a 32^3^ voxel overlap to guarantee that the core patches within all patches collectively cover the entirety of the original cryo-EM map seamlessly.

Our approach involves the use of two distinct patch sizes, 64 and 128, for dividng the cryo-EM maps into patches. Consequently, we obtain two sets of predicted atomic classifications. Each voxel is assigned a 19-dimensional vector representing the probabilities of the 19 atom classes, and the atom class with the highest probability is chosen as the final predicted atom class for that voxel. After neural network prediction, we have two sets of predicted atoms, referred to as “atom-ps64” and “atom-ps128”, respectively.

### 4.5 Atom clustering

Ideally, the voxel corresponding to a truth-ground atom and its directly neighboring six voxels should be predicted to contain atoms of the same class as the truth-ground atom, consistent with the training data. However, when applying the trained neural network to both the test and training datasets, much more voxels surrounding the truth-ground atom are predicted to contain atoms. Therefore, we need to cluster the predicted atoms into representative groups as follows:

(1) For each predicted RNA atom, we calculate the number of atoms belonging to the same RNA atom class within a 3^3^ voxel box centered at that atom. When counting neighboring atoms, we treat base atom pairs like N9-N1, C4-C2, and C8-C6 as the same atom class.
(2) We employ the “peak local max” function from the scikit-image Python package [50] to identify local peaks for each RNA atom class, which are defined as voxels with the maximum number of neighboring atoms in our approach. Two peak voxels have to be separated by at least five voxels, corresponding to a distance of 2.5 Å in 3D space, given that each voxel has a size of 0.5 Å after preprocessing the cryo-EM map. Additionally, a peak voxel must have at least five neighboring atoms to exclude the predicted “orphan” atoms, which could be artifacts caused by noise.
(3) The identified peak voxels are labeled with the atom classes that are employed to recognize these local peaks. For a found peak in the base atom class, such as N9 and N1, which are treated as the same atom class when counting neighboring atoms, we separately count the numbers of atom N9 and N1 within a 5^3^ voxel box centered at the found peak. The found peak is then assigned as the atom class N9 or N1 based on which class has more atoms within this box. Similarly, the same process is applied to the other two base atom pairs C4-C2, and C8-C6. This atom clustering approach allows us to reduce the redundant atoms into representative ones.

### 4.6 Nucleotide assembly from atoms

We synthesize nucleotides by assembling previously clustered atoms, guided by their pairwise atomic distances. The process involves the following steps:

(1) Seed selection: Initially, we designate the C4’ atoms as seeds for nucleotide growth.
(2) Atom addition: Subsequently, atoms are added one by one to the growing nucleotide, considering the distance between the candidate atom and the reference atom. For instance, we add a candidate C5’ atom into the nucleotide if it is the nearest C5’ atom to the reference C4’ atom within a 3.0 Å distance threshold. Various candidate atoms are associated with distinct reference atoms and corresponding distance thresholds. The specific candidate atoms, their corresponding reference atoms, and distance thresholds are detailed in Table 1. In some cases, a candidate atom may have two reference atoms, with the second one used if the first is absent in the nucleotide. If no reference atom exists for a candidate atom, the candidate atom is omitted from the nucleotide, and it will be reconstructed later according to nucleotide templates.
(3) Bond length and nucleotide size check: We inspect covalent bond lengths in a standard nucleotide and remove atoms with bond lengths exceeding 3.0 Å. Nucleotides with fewer than five atoms after the atom addition step are removed.
(4) Nucleotide type determination: The type of a nucleotide is determined by the number of specific base atoms. If a nucleotide contains more N9, C4, and C8 atoms than N1, C2, and C6 atoms, it is classified as an adenine nucleotide (abbreviated as A) and N1, C2, and C6 atoms will be removed. Conversely, it is designated as a cytosine nucleotide (abbreviated as C) and atoms N9, C4, and C8 will be removed. If a nucleotide lacks base atoms, it is classified as an unknown nucleotide type and denoted as X.
(5) Backbone and base reconstruction: The missing RNA backbone and base atoms are reconstructed using pools of RNA backbone and sugar-base templates, respectively.

**Table 1.** Candidate and reference atoms used in atom addition.

| Candidate atom | Reference atom | Dis. threshold (Å) |
| --- | --- | --- |
| C5' | C4' | 3.0 |
| C3' | C4' | 3.0 |
| O4' | C4' | 3.0 |
| O5' | C5' | 3.0 |
| O5' | C4' | 3.5 |
| C2' | C3' | 3.0 |
| C2' | C4' | 3.5 |
| O3' | C3' | 3.0 |
| O3' | C4' | 3.5 |
| C1' | O4' | 3.0 |
| C1' | C4' | 3.5 |
| O2' | C2' | 3.0 |
| O2' | C4' | 3.8 |
| P | O5' | 3.0 |
| P | C5' | 4.0 |
| OP1 | P | 3.0 |
| OP1 | O5' | 4.0 |
| OP2 | P | 3.0 |
| OP2 | O5' | 4.0 |
| N9 | C1' | 3.0 |
| N9 | C4' | 5.5 |
| N1 | C1' | 3.0 |
| N1 | C4' | 5.5 |
| C8 | C1' | 4.0 |
| C8 | C4' | 5.5 |
| C6' | C1' | 4.0 |
| C6' | C4' | 5.5 |
| C4 | C1' | 4.0 |
| C4 | C4' | 6.5 |
| C2 | C1' | 4.0 |
| C2 | C4' | 6.5 |

Following nucleotide assembly, we obtain two sets of nucleotides, designated as “nt-ps64” and “nt-ps128”, aligning with the two respective sets of atoms, “atom-ps64” and “atom-ps128”.

### 4.7 Linking nucleotides into short chains

Upon identifying all nucleotides through atom clustering and assembly, we proceed to synthesize short nucleotide chains by covalently linking those that are in close proximity. This process comprises several key steps:

(1) Determining neighboring nucleotides: We first define neighboring (covalently bonded) nucleotides based on specific distance criteria. Nucleotides *i* and *j* are considered neighboring if they meet any of the following four distance criteria: (a) If the distance between the O3’ atom in nucleotide *i* and the P atom in nucleotide *j* is less than 3.0 Å. (b) If the distance between the C3’ atom in nucleotide *i* and the P atom n nucleotide *j* is less than a predetermined threshold. This threshold is set at 4.5 Å when both the RMSDs of the sugar groups, consisting of atoms C5’, C4’, C3’, C2’, C1’, and O4’, in nucleotides *i* and *j* relative to a standard sugar group, are less than 0.6 Å. Otherwise, it is set at 4.0 Å. (c) If the P-O5’ bond length in nucleotide *j* mea-sures less than 2.6 Å and the distance between the O3’ atom in nucleotide *i* and the O5’ atom in nucleotide *j* is less than 3.5 Å. (d) If the P-O5’ bond length in nucleotide *j* measures less than 2.6 Å and the distance between the C3’ atom in nucleotide *i* and the O5’ atom in nucleotide *j* is less than 4.0 Å. In this context, we establish that nucleotide *i* serves as the upstream neighbor of nucleotide *j*, while nucleotide *j* is considered the downstream neighbor of nucleotide *i*. It is worth noting that a nucleotide may have multiple upstream and/or downstream neighbors.
(2) Short chain growth: Following neighbor identification, we extend nucleotide chains bidirectionally. Starting from a seeding nucleotide, we incorporate its upstream and downstream neighbors into a short chain. This process continues by adding the upstream neighbor of the first nucleotide and the downstream neighbor of the last nucleotide, extending the short chain until the first and last nucleotides have no neighboring nucleotides. Subsequently, new short chains are initiated from seeding nucleotides not already part of existing short chains. It’s worth emphasizing that when a nucleotide has multiple neighboring nucleotides, all possible growth pathways for short chains are considered. If there are *N* sets of neighboring nucleotides, *N* sets of short chains are generated.
(3) Short chain pruning: To streamline subsequent chain extension and sequence alignment, we eliminate short chains comprising fewer than four nucleotides.
(4) Helical region chain splitting: In situations where it is possible for a hairpin loop to form following chain splitting, we divide the chains within the helical region. Specifically, if the distance between the C4’ atoms in nucleotide *n* in the helical region of chain *i* and the head or tail nucleotide in another chain *j* is less than 10 Å, and if chain *j* and sub-chains *i*1 or *i*2 after splitting have the potential to create a hairpin loop, we split chain *i* at nucleotide *n*. This is illustrated in Fig. 8. In Fig. 8a, chains A and B exhibit the formation of kissing hairpins within an RNA native structure. Moving to Fig. 8b, our approach yields three short chains, denoted as C, D, and E. It is evident that certain segments of chains A and B, originating from the native structure, have been incorrectly merged into the red-colored chain E in Fig. 8b. In Fig. 8b, the C4’ atoms in the tail nucleotide of chain C and the middle nucleotide in chain E are in close proximity less than 10 Å, marked by the dashed arrow. After splitting at the arrowhead, as depicted in Fig. 8c, chain C and the newly formed sub-chain F can come together to create a hairpin loop. For RNAs containing kissing hairpin loops, the chain splitting process provides the opportunity to generate correct chain structures.

**Fig. 8.**
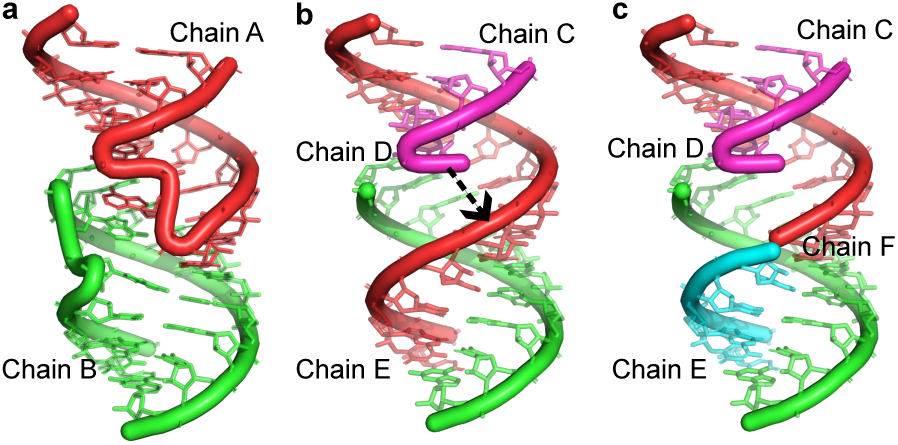
Short chain splitting. **a,** The native RNA structure depicting the kissing hairpin, with chains A and B highlighted in red and green, respectively. **b,** Predicted structures featuring three short chains: C in magenta, D in green, and E in red. The arrowhead marks the potential splitting site in chain E. **c,** Following the chain split shown in panel b, chain E is divided into sub-chains E and F. The resulting sub-chain F and chain C have the potential to form a hairpin loop.

### 4.8 Extending short chains into complete long chains

Within each of the *N* sets of short chains, we implement a comprehensive approach to establish connectivity among short chains, ultimately forming complete long chains. This multi-step procedure includes:

(1) Identifying neighboring short chains: We identify neighboring short chains when either of the following two criteria is satisfied: (a) If the distance between the C4’ atom in the last nucleotide of short chain *i* and the C4’ atom in the first nucleotide of short chain *j* (referred to as chain dis*_ij_*) is less than 20 Å. (b) If the RMSD of the C4’ atoms in the last two nucleotides of short chain *i* and the C4’ atoms in the first two nucleotides of short chain *j*, with respect to the four C4’ atoms in a standard single-stranded A-helix, is less than 1.5 Å. It is important to note that the first two C4’ atoms n the reference A-helix are continuous in sequence, as well as the last two C4’ atoms, but the second and third C4’ atoms may exhibit gaps of zero to eight nucleotides. This RMSD value is denoted as chain RMSD*_ij_*. We employ this RMSD criterion to infer that these two short chains are likely connected by a single-stranded helix, even if the helical structure is not present in the predicted structures. We designate short chain *i* as the upstream neighbor of short chain *j*, and short chain *j* as the downstream neighbor of short chain *i*.
(2) Filtering neighboring chains: When a set of short chains comprises more than eight individual chains, we filter short chain *i*’s downstream neighbors when multiple downstream neighbors are present. Specifically, we exclude short chain *j* from short chain *i*’s downstream neighbors if the chain dis*_ij_* exceeds 10 Å, and chain dis*_ij_* is greater by 6 Å compared to the minimum chain dis*_ik_* value, where *k* represents all the downstream neighbors of short chain *i*, and the chain RMSD*_ij_* exceeds 1.5 Å. Accordingly, short chain *i* no longer acts as the upstream neighbor of short chain *j* after filtering.
(3) Forming complete long chains: In this phase, we form complete long chains by weaving the short chains together to create a comprehensive structure. This process commences by extending a long chain from a chosen seed short chain, considering the neighboring short chains determined above. Four distinct options are employed to extend the long chain: (a) Only the upstream neighbor of the first short chain is added to the long chain. (b) Conversely, only the downstream neighbor of the last short chain is integrated into the long chain. (c) A combination of both neighboring chains is included, following options (a) and (b). (d) Instead of extending the current long chain, we select a new short chain not in the existing long chain as the seed to develop a new long chain. This choice is particularly valuable when dealing with RNA structures that exhibit multiple sequence chains, which may not be spatially linked in 3D space. It is noteworthy that if a long chain lacks both upstream and downstream neighbors, we also utilize option (d) to initiate a new long chain. After this step, a variety of complete chains, comprising one or multiple long chains, can be generated.
(4) Filtering complete long chains: To prevent redundancy, we employ a filtering process to remove duplicate complete chains. This is necessary since the four extension options described earlier can lead to the generation of identical complete chains. Furthermore, we discard complete chains in which the number of long chains surpasses the native RNA’s chain count by more than four. This is executed to ensure that the resulting complete long chains maintain a reasonable level of continuity, avoiding excessive fragmentation compared to the native chain structure.

In summary, a set of short chains is employed for long chain extension to construct complete long chains, and this process is repeated for all sets of short chains.

### 4.9 Sequence assignment through global sequence alignment

To obtain the accurate sequence and chain information for the aforementioned complete long chains, global sequence alignment is performed between the query sequences from the complete long chains and the reference sequences from the provided native RNA chains. Subsequently, the native sequence information is assigned to the complete long chains based on the top best alignments. This process consists of several sequential steps:

(1) Conversion of complete long chains into query sequences: Initially, we systematically transform each complete long chain, composed of multiple short chains, into a query sequence. This transformation involves converting each short chain into a sub-sequence containing the predicted nucleotides A, C, and X. Subsequently, we link these sub-sequences from individual short chains together, using the symbol “N” to signify short chain separation. Moreover, to ensure that the short chains remain separated by a minimal number of nucleotides, we introduce placeholder symbols “P” between them immediately following the symbol “N”. The determination of the number of “P” symbols, referred to as *n*P, is guided by the distance chain dis*_ij_* between the C4’ atom n the last nucleotide of short chain *i* and the C4’ atom in the first nucleotide of the downstream neighboring short chain *j* in the complete chain. The calculation of *n*P s performed using the formula *n*P = min(max([chain dis*_ij_/*6] − 1, 0), 3). In this context, the value 6 represents an approximate atomic distance between two C4’ atoms n adjacent nucleotides within an A-helix, while 3 denotes the maximum allowable number of “P” symbols inserted between short chains. After conversion, these query sequences include letters A, C, X, N, and P.
(2) Preparation of reference sequences: Subsequently, we generate reference sequences that correspond to the provided native RNA chains. In cases where the RNA structure comprises multiple chains, we systematically enumerate all feasible chain permutations. By connecting these permutations with the symbol “T”, signifyng chain termination, we create comprehensive reference sequences. These reference sequences encompass letters A, C, G, U, and potentially T.
(3) Global sequence alignment using a modified Gotoh algorithm: To achieve global sequence alignment, we employ a modified Gotoh algorithm [33]. This specialized alignment technique introduces distinct gap opening and extending costs. In the reference sequence, both gap opening and extending costs are set to -2. However, in the query sequence, the gap opening and extending costs are variable based on their specific position. If gaps are opened or extended immediately before or after the “N” symbol within the query sequence, they are assigned costs of -2. On the other hand, if gaps occur at other positions, they are penalized more severely with opening and extending costs of -5 and -4, respectively. This differentiation is specifically designed to discourage the presence of gaps within the short chains, which are covalently bonded. The scoring matrix employed for our sequence alignment protocol is comprehensively detailed in Table 2.
(4) Identification of top alignments: As part of our effort to expedite the alignment process and identify the most suitable alignments, we only calculate the raw best alignment score for each query-reference sequence pair without tracing back the detailed alignments. This calculation is efficiently executed in C++. For the top 50 query-reference sequence pairs, we conduct comprehensive sequence alignments and subsequently retrieve detailed alignment results.
(5) Assignment of native sequence: Building upon the top 10 alignment results, we proceed to assign the native sequence information to the predicted complete chain structures. This assignment process involves making essential adjustments to the base atoms and nucleotide names, as well as rearranging and renumbering the nucleotides in the sequence.

**Table 2.** Scoring matrix for global sequence alignment.

|  | A | C | G | U | T |
| --- | --- | --- | --- | --- | --- |
| A | 1 | -1 | 1 | -1 | -5 |
| C | -1 | 1 | -1 | 1 | -5 |
| X | -1 | -1 | -1 | -1 | -5 |
| N | -3 | -3 | -3 | -3 | 5 |
| P | -2 | -2 | -2 | -2 | -2 |

By following these procedures, we can obtain up to 10 predicted all-atom structures for each of the two sets of nucleotides “nt-ps64” and “nt-ps128”. Additionally, we identify the top 10 predicted structures by assessing alignment scores for both patch sizes. In total, our comprehensive approach yields up to 30 predicted structures, categorized nto three groups in terms of patch size, denoted as “RNA-ps64”, “RNA-ps128”, and “RNA-psBoth”.

### 4.10 Energy minimization for refinement

For the rebuilt all-atom RNA structures, we employ the software QRNAS [34] to conduct energy minimization, addressing broken bonds and eliminating steric clashes between atoms. This local refinement process minimally alters the structural conformation, as indicated by a negligible change in RMSD less than 0.05 Å.

## 5 Data availability

The cryo-EM maps and RNAs used for training and test are listed in Supplementary Table 1 and 2, respectively. The experimental cryo-EM maps listed in the tables can be downloaded from the EMDB (https://www.emdataresource.org/). The native RNA structures listed in the tables can be downloaded from the RCSB PDB (https://www.rcsb.org/). The 51 test RNA structures predicted by Deep-Tracer, CryoREAD, and DeepCryoRNA can be found at https://github.com/Vfold-RNA/DeepCryoRNA/releases/tag/v1.0. For DeepCryoRNA, the complete working directories containing all the inputs and outputs for the 51 test RNAs are also included.

## 6 Code availability

The source code and tutorial for DeepCryoRNA is available at https://github.com/Vfold-RNA/DeepCryoRNA.

## Supplementary information

Supplementary material is attached.

## Acknowledgments

This work was supported by the National Institutes of Health under Grant R35-GM134919 to S-J. C.

