## Supplementary material for "DeepCryoRNA: deep learning-based RNA structure reconstruction from cryo-EM maps": SI

**Supplementary Table 1. Training dataset.**

| PDB | EMD | Map resolution | RNA length | Contour |
|---|---|---|---|---|
| 6XZ7 | 10655 | 2.1 | 3126 | 0.09 |
| 7OF0 | 12865 | 2.2 | 1432 | 0.004 |
| 6AZ3 | 7025 | 2.5 | 3439 | 0.065 |
| 7N0D | 24104 | 2.5 | 76 | 0.433 |
| 6TH6 | 10503 | 2.55 | 4441 | 0.05 |
| 7O9M | 12764 | 2.6 | 1513 | 0.15 |
| 7KGB | 22865 | 2.7 | 4815 | 0.02 |
| 7MQA | 23938 | 2.7 | 1763 | 3.28 |
| 7R81 | 24307 | 2.7 | 5339 | 0.5 |
| 6SGC | 10181 | 2.8 | 5708 | 0.05 |
| 7C7A | 30297 | 2.8 | 339 | 0.006 |
| 5T2A | 8343 | 2.9 | 5278 | 0.01 |
| 6W5C | 21541 | 2.9 | 50 | 0.0428 |
| 6YYT | 11007 | 2.9 | 54 | 3 |
| 6RJA | 4901 | 3 | 156 | 0.0625 |
| 6XU8 | 10624 | 3 | 5912 | 0.035 |
| 7ONU | 13002 | 3 | 66 | 5 |
| 7PLA | 13486 | 3.04 | 217 | 0.055 |
| 7DMQ | 30767 | 3.06 | 88 | 0.0038 |
| 6XMF | 22257 | 3.1 | 109 | 0.0254 |
| 6ZMO | 11299 | 3.1 | 5951 | 0.05 |
| 7VG2 | 31963 | 3.1 | 71 | 0.0436 |
| 6ZJ3 | 11232 | 3.15 | 5716 | 0.035 |
| 6D90 | 7834 | 3.2 | 5842 | 0.025 |
| 5M1J | 4140 | 3.3 | 5429 | 3.5 |
| 6E9F | 9014 | 3.3 | 79 | 0.2 |
| 6SV4 | 10315 | 3.3 | 15871 | 0.02 |
| 6SXO | 10344 | 3.3 | 326 | 0.01 |
| 7ELH | 31183 | 3.3 | 310 | 0.006 |
| 7ASE | 11893 | 3.33 | 2225 | 0.0193 |
| 7RE3 | 24432 | 3.33 | 142 | 0.4 |
| 7EGQ | 31138 | 3.35 | 116 | 0.25 |
| 6NM9 | 9398 | 3.38 | 50 | 0.0177 |
| 5MMM | 3533 | 3.4 | 4585 | 0.06 |
| 5ZWN | 6973 | 3.4 | 502 | 0.045 |
| 6MCB | 9066 | 3.4 | 115 | 0.28 |
| 6RM3 | 4935 | 3.4 | 3662 | 0.0199 |
| 7BGB | 12177 | 3.4 | 92 | 0.022 |
| 6XN7 | 22269 | 3.47 | 74 | 0.0185 |
| 6AH3 | 9622 | 3.48 | 449 | 0.025 |
| 6ZVK | 11459 | 3.49 | 6218 | 0.027 |
| 6RXU | 10052 | 3.5 | 1805 | 0.03 |

| | | | | |
|---|---|---|---|---|
| 6XU6 | 10622 | 3.5 | 5916 | 0.025 |
| 7ASA | 11891 | 3.5 | 116 | 0.015 |
| 5T2C | 8345 | 3.6 | 5869 | 0.03 |
| 5TC1 | 8397 | 3.6 | 242 | 3 |
| 6ME0 | 9105 | 3.6 | 829 | 0.023 |
| 7KRO | 23008 | 3.6 | 80 | 0.4 |
| 6AHU | 9627 | 3.66 | 413 | 0.0256 |
| 6J6Q | 692 | 3.7 | 549 | 0.0116 |
| 6OM7 | 597 | 3.7 | 5725 | 0.022 |
| 6V5B | 21051 | 3.7 | 78 | 0.6 |
| 6Z1P | 11032 | 3.7 | 4096 | 0.03 |
| 5VT0 | 8732 | 3.78 | 112 | 0.014 |
| 5A8L | 3099 | 3.8 | 347 | 0.0008 |
| 6O1O | 454 | 3.8 | 66 | 0.9 |
| 6WDA | 21629 | 3.8 | 4810 | 1 |
| 7BG9 | 12174 | 3.8 | 256 | 0.014 |
| 3J7R | 2644 | 3.9 | 5898 | 0.065 |
| 5H0R | 9564 | 3.9 | 84 | 12 |
| 6LXD | 30005 | 3.9 | 72 | 0.02 |
| 7ANE | 11829 | 3.9 | 1688 | 0.045 |
| 7L48 | 23157 | 3.9 | 144 | 0.0146 |
| 7NSJ | 12569 | 3.9 | 2719 | 0.025 |
| 7OBQ | 12799 | 3.9 | 165 | 0.01 |
| 5ODV | 3785 | 4 | 120 | 0.021 |
| 6T83 | 10398 | 4 | 10549 | 0.02 |
| 6ZS9 | 11390 | 4 | 2487 | 0.02 |
| 5KPV | 8280 | 4.1 | 4812 | 0.1 |
| 7JQQ | 22441 | 4.1 | 585 | 0.0114 |
| 6GZZ | 105 | 4.13 | 9080 | 3.1 |
| 5JUS | 6645 | 4.2 | 5567 | 0.015 |
| 6NY1 | 8994 | 4.2 | 108 | 0.3 |
| 6WB0 | 21583 | 4.2 | 73 | 0.02 |
| 4V8Y | 2421 | 4.3 | 5354 | 0.25 |
| 6K0B | 9900 | 4.3 | 682 | 0.02 |
| 6ND4 | 441 | 4.3 | 551 | 0.0203 |
| 7JL2 | 22370 | 4.3 | 88 | 0.02 |
| 6RXZ | 10056 | 4.4 | 1717 | 0.03 |
| 5WYK | 6696 | 4.5 | 1619 | 0.09 |
| 6G4W | 4349 | 4.5 | 1563 | 0.04 |
| 6GZX | 104 | 4.57 | 9080 | 2.3 |
| 6CB1 | 7445 | 4.6 | 1615 | 0.018 |
| 7AJT | 11807 | 4.6 | 2025 | 0.01 |
| 7LHD | 23336 | 4.6 | 4217 | 0.018 |
| 3JBN | 6456 | 4.7 | 5146 | 0.03 |
| 5ZAL | 6905 | 4.7 | 58 | 0.04 |
| 6D6V | 7821 | 4.8 | 159 | 0.055 |
| 6LQV | 955 | 4.8 | 1296 | 0.022 |

| | | | | |
|---|---|---|---|---|
| 6XMG | 22258 | 4.8 | 154 | 0.0125 |
| 6P7N | 20267 | 4.9 | 50 | 0.08 |
| 6ZQF | 11362 | 4.9 | 1447 | 0.01 |
| 7ELE | 31182 | 4.9 | 83 | 0.006 |
| 5MRF | 3553 | 4.97 | 4286 | 0.0123 |
| 4V6X | 5592 | 5 | 6512 | 0.7 |
| 5AA0 | 6397 | 5 | 4753 | 0.0392 |
| 5TZS | 8473 | 5.1 | 629 | 0.018 |
| 5Y36 | 8236 | 5.2 | 99 | 0.0316 |
| 6T34 | 10373 | 5.2 | 95 | 0.03 |
| 6I7O | 4427 | 5.3 | 10687 | 0.013 |
| 4V7E | 1780 | 5.5 | 5519 | 0.11 |
| 4V8M | 2239 | 5.57 | 6500 | 108000 |
| 5MYJ | 3581 | 5.6 | 4547 | 0.035 |
| 6AH0 | 9621 | 5.7 | 452 | 0.015 |
| 7OI6 | 12919 | 5.7 | 1264 | 0.02 |
| 3JBO | 6452 | 5.8 | 5144 | 0.02 |
| 5LQW | 4099 | 5.8 | 378 | 0.11 |
| 5MQF | 3545 | 5.9 | 387 | 0.016 |
| 7OQE | 13033 | 5.9 | 734 | 0.0125 |
| 4V6W | 5591 | 6 | 6111 | 0.49 |
| 7SCQ | 25041 | 6 | 169 | 0.0318 |
| 3J6X | 5942 | 6.1 | 5368 | 0.815 |
| 3J77 | 5976 | 6.2 | 5433 | 0.12 |
| 3J78 | 5977 | 6.3 | 5525 | 0.12 |
| 4V68 | 5030 | 6.4 | 4695 | 3.5 |
| 2WWB | 1652 | 6.48 | 136 | 0.2 |
| 5Z57 | 6890 | 6.5 | 390 | 0.0323 |
| 6HCQ | 197 | 6.5 | 5817 | 0.07 |
| 7PJX | 13463 | 6.5 | 4682 | 1.5 |
| 4V6U | 2009 | 6.6 | 4822 | 0.13 |
| 4V8Z | 2422 | 6.6 | 5272 | 0.15 |
| 3JBP | 6454 | 6.7 | 5144 | 0.03 |
| 4V69 | 5036 | 6.7 | 4789 | 90 |
| 6FXC | 3637 | 6.76 | 9118 | 0.023 |
| 4V7B | 5775 | 6.8 | 4677 | 3 |
| 6HCM | 195 | 6.8 | 5736 | 0.07 |
| 4UJE | 2620 | 6.9 | 5813 | 3.5 |
| 3JD5 | 5941 | 7 | 952 | 0.00017 |
| 4UE4 | 2843 | 7 | 266 | 100 |
| 6RXT | 10051 | 7 | 1382 | 0.03 |
| 6XII | 22192 | 7 | 4706 | 0.00722 |

**Supplementary Table 2. Testing dataset.**

| PDB | EMD | Map resolution | RNA length | Contour |
|---|---|---|---|---|
| 7YGA | 33813 | 2.35 | 407 | 0.3 |
| 7YGB | 33814 | 2.62 | 407 | 0.25 |
| 7YGC | 33815 | 2.65 | 407 | 0.14 |
| 7YG9 | 33812 | 2.68 | 404 | 0.01 |
| 7XD5 | 33136 | 2.84 | 406 | 0.01 |
| 7XD6 | 33137 | 2.84 | 459 | 0.01 |
| 7R6L | 24281 | 2.85 | 366 | 0.393 |
| 7YG8 | 33811 | 2.97 | 395 | 0.18 |
| 7YCI | 33740 | 2.98 | 395 | 0.1 |
| 8I7N | 35223 | 2.98 | 418 | 0.02 |
| 8SA3 | 40263 | 3 | 420 | 0.363 |
| 7XSN | 33428 | 3.01 | 387 | 0.27 |
| 7XD7 | 33138 | 3.02 | 405 | 0.01 |
| 7EZ2 | 31386 | 3.05 | 406 | 0.023 |
| 7YCH | 33739 | 3.09 | 404 | 0.1 |
| 8SA2 | 40262 | 3.1 | 420 | 0.361 |
| 8SA4 | 40264 | 3.1 | 267 | 0.377 |
| 7EZ0 | 31385 | 3.14 | 387 | 0.007 |
| 7YCG | 33738 | 3.18 | 404 | 0.1 |
| 7YGD | 33816 | 3.41 | 389 | 0.2 |
| 8SA5 | 40265 | 3.5 | 345 | 0.38 |
| 8HD7 | 34671 | 3.52 | 419 | 0.006 |
| 7XSK | 33425 | 3.53 | 388 | 0.27 |
| 7R6M | 24282 | 3.68 | 362 | 0.237 |
| 6UES | 20755 | 3.7 | 119 | 9 |
| 8HD6 | 34670 | 3.73 | 419 | 0.006 |
| 7XSL | 33426 | 3.84 | 388 | 0.22 |
| 7XD4 | 33135 | 3.89 | 423 | 0.017 |
| 7UVT | 26816 | 3.9 | 386 | 0.075 |
| 7XSM | 33427 | 4.01 | 388 | 0.22 |
| 7XD3 | 33134 | 4.05 | 423 | 0.017 |
| 7PTQ | 13633 | 4.08 | 544 | 0.15 |
| 6UET | 20756 | 4.1 | 119 | 5 |
| 7R6N | 24283 | 4.1 | 354 | 0.225 |
| 7YC8 | 33736 | 4.14 | 388 | 0.1 |
| 7SAM | 24952 | 4.3 | 169 | 0.0264 |
| 7ZJ4 | 14740 | 4.43 | 374 | 0.14 |
| 5G2Y | 3332 | 4.5 | 630 | 0.04 |
| 7ZJ5 | 14741 | 4.55 | 374 | 0.105 |
| 6WLQ | 21838 | 4.7 | 119 | 0.7 |
| 6WLR | 21839 | 4.8 | 119 | 1.2 |

| | | | | |
|---|---|---|---|---|
| 6WLT | 21841 | 4.8 | 231 | 0.8 |
| 6POM | 20416 | 4.9 | 230 | 0.046 |
| 7PTL | 13630 | 4.9 | 720 | 0.4 |
| 7QDU | 13926 | 5.14 | 552 | 0.138 |
| 7PTK | 13628 | 5.18 | 720 | 0.1 |
| 8SA6 | 40266 | 5.3 | 350 | 0.169 |
| 8BTZ | 16244 | 5.39 | 238 | 0.124 |
| 7LJY | 23401 | 5.6 | 101 | 0.459 |
| 6WLU | 21842 | 5.7 | 231 | 0.2 |
| 7PTS | 13636 | 5.71 | 558 | 0.366 |

**Supplementary Table 3. The RMSDs achieved by five methods for the 51 testing RNAs.** The unit of RMSD is angstrom (Å).

| PDB | DeepCryoRNA _ps64 | DeepCryoRNA _ps128 | DeepCryoRNA _psBoth | DeepTracer | CryoREAD |
|---|---|---|---|---|---|
| 7YGA | 2 | 2.1 | 2 | 32.1 | 41.9 |
| 7YGB | 2.4 | 2.1 | 2.4 | 43 | 43.8 |
| 7YGC | 3.1 | 2 | 3.1 | 40 | 41.6 |
| 7YG9 | 2.6 | 2.6 | 2.6 | NaN | 46.6 |
| 7XD5 | 1.9 | 2.1 | 2.1 | 41.1 | 41.1 |
| 7XD6 | 5.9 | 6 | 6 | 45.9 | 46.2 |
| 7R6L | 2.8 | 1.6 | 1.6 | 50.4 | 38 |
| 7YG8 | 3.4 | 3.4 | 3.4 | NaN | 43.4 |
| 7YCI | 2.4 | 2.3 | 2.4 | NaN | 43.9 |
| 8I7N | 2.7 | 4.5 | 4.5 | 43.5 | 46.1 |
| 8SA3 | 26 | 5 | 5 | NaN | 57.2 |
| 7XSN | 1.8 | 2.1 | 2.1 | NaN | 40.8 |
| 7XD7 | 7.6 | 5.8 | 5.8 | 38.1 | 42.6 |
| 7EZ2 | 2 | 2.1 | 2 | 46.9 | 45.7 |
| 7YCH | 2.3 | 2.1 | 2.1 | NaN | 42.4 |
| 8SA2 | 44.8 | 6 | 6 | NaN | 56.6 |
| 8SA4 | 6.4 | 3.4 | 3.4 | NaN | 41.3 |
| 7EZ0 | 2.8 | 2.2 | 2.2 | NaN | 40.8 |
| 7YCG | 2.5 | 2.5 | 2.5 | NaN | 39.9 |
| 7YGD | 3 | 3.9 | 3 | NaN | 43.3 |
| 8SA5 | 42.2 | 18.7 | 18.7 | NaN | 43.5 |
| 8HD7 | 9.9 | 6.1 | 6.1 | NaN | 41.6 |
| 7XSK | 3.5 | 11.2 | 3.5 | NaN | 41.7 |
| 7R6M | 7.5 | 3.9 | 3.9 | NaN | 38.3 |
| 6UES | 2.6 | 2.4 | 2.4 | NaN | 18.4 |
| 8HD6 | 10.5 | 3.7 | 3.7 | NaN | 45.3 |
| 7XSL | 4.5 | 6.2 | 4.5 | NaN | 42.4 |
| 7XD4 | 34.5 | 5.1 | 5.1 | NaN | 46.6 |
| 7UVT | 5.5 | 5.5 | 5.5 | NaN | 41.8 |
| 7XSM | 30 | 13.3 | 30 | NaN | 40.8 |
| 7XD3 | 10.8 | 2.5 | 2.5 | NaN | 48.9 |
| 7PTQ | 33.1 | 5.9 | 5.9 | NaN | 55.7 |
| 6UET | 2.9 | 3 | 3 | NaN | 24 |
| 7R6N | 6.4 | 2.6 | 2.6 | 47.9 | 41.4 |
| 7YC8 | 20.9 | 26.5 | 20.9 | NaN | 43.7 |
| 7SAM | 16.5 | 14.9 | 16.5 | NaN | 30.2 |
| 7ZJ4 | 22.6 | 18.2 | 18.2 | NaN | 61.4 |
| 5G2Y | NaN | 52.3 | 52.4 | NaN | 64.6 |
| 7ZJ5 | 38.5 | 46.3 | 46.3 | NaN | 53.4 |
| 6WLQ | 6.8 | 4 | 6.8 | NaN | 29.2 |
| 6WLR | 26.3 | 5.8 | 5.7 | NaN | 26.3 |
| 6WLT | 5.9 | 10.7 | 5.9 | NaN | 35.9 |

| | | | | | |
|---|---|---|---|---|---|
| 6POM | 9.2 | 6.9 | 6.9 | NaN | 38.4 |
| 7PTL | NaN | NaN | NaN | NaN | 58.8 |
| 7QDU | 64.8 | NaN | 64.8 | NaN | 65.5 |
| 7PTK | NaN | NaN | NaN | NaN | 60.2 |
| 8SA6 | 70 | 38.8 | 70 | NaN | 73.6 |
| 8BTZ | 23 | 28.7 | 28.7 | NaN | 39.8 |
| 7LJY | 35.7 | 14.9 | 14.9 | NaN | 32.9 |
| 6WLU | 33.5 | 30.7 | 33.5 | NaN | 39.7 |
| 7PTS | 40.7 | NaN | 40.7 | NaN | 52.6 |
